# The structural basis of the multi-step allosteric activation of Aurora B kinase

**DOI:** 10.1101/2022.11.23.517677

**Authors:** Dario Segura-Peña, Oda Hovet, Hemanga Gogoi, Jennine M. Dawicki-McKenna, Stine Malene Hansen Wøien, Manuel Carrer, Ben E. Black, Michele Cascella, Nikolina Sekulić

## Abstract

Aurora B, together with IN-box, the C-terminal part of INCENP, forms an enzymatic complex that ensures faithful cell division. The [Aurora B/IN-box] complex is activated by autophosphorylation in the Aurora B activation loop and in IN-box, but it is not clear how these phosphorylations activate the enzyme. We used a combination of experimental and computational studies to investigate the effects of phosphorylation on the molecular dynamics and structure of [Aurora B/IN-box]. In addition, we generated partially phosphorylated intermediates to analyze the contribution of each phosphorylation independently. We found that the dynamics of Aurora and IN-box are interconnected, and IN-box plays both positive and negative regulatory roles depending on the phosphorylation status of the enzyme complex. Phosphorylation in the activation loop of Aurora B occurs intramolecularly and prepares the enzyme complex for activation, but two phosphorylated sites are synergistically responsible for full enzyme activity.

## Introduction

The chromosome passenger complex (CPC) is a complex of four proteins involved in the regulation of key mitotic events: correction of chromosome-microtubule attachment errors, assembly and stability of the spindle assembly checkpoint, and assembly and regulation of the contractile apparatus that drives cytokinesis (Carmena et al., 2012; Vader et al., 2006). The CPC is organized into two modules. The localization module consists of survivin, borealin, and the N-terminal part of INCENP, whereas the enzymatic module comprises Aurora B kinase and the C-terminal part of INCENP, called the IN-box. The enzymatic activity of Aurora B is essential for the regulatory functions of CPC (Kelly and Funabiki, 2009; Krenn and Musacchio, 2015; Meraldi et al., 2004) so Aurora B has been identified as an attractive target for cancer chemotherapy (Portella et al., 2011), with several inhibitors patented in the last decades (Jing and Chen, 2021).

The enzymatic activity of the [Aurora B/IN-box] is tightly regulated by autophosphorylation. The activity of the complex increases by at least two orders of magnitude after the complex phosphorylates itself (Zaytsev et al., 2016). There are two well-defined autophosphorylation sites in the [Aurora B/IN-box] enzyme complex. One is a threonine in the Aurora B activation loop, and the second is located at the C-terminus of the IN-box and consists of two consecutive serine residues in the TSS motif (Bishop and Schumacher, 2002; Honda et al., 2003; Sessa et al., 2005).

Until recently, the molecular understanding of Aurora B autoactivation was based on a model derived from the structure of the partially phosphorylated enzyme, in which the threonine in the Aurora B activation loop was phosphorylated but the TSS motif of INCENP was either absent (Sessa et al., 2005; Sessa and Villa, 2014) or unstructured (Elkins et al., 2012). Recently, a structural study of the [Aurora C/IN-box] complex, which is specific for meiosis but otherwise very similar to the [Aurora B/IN-box] complex, provided valuable insight into the conformation the fully phosphorylated enzyme adopts (Abdul Azeez et al., 2019). The crystal structure shows that the activation loop of Aurora C and the IN-box TSS motif interact in the phosphorylated state and stabilize the extended (active) conformation of the activation loop. Although the enzyme complex was crystallized with an inhibitor bound to the ATP-binding site, it is likely that the observed interaction between the IN-box and the activation loop represents a conformation corresponding to the active state of the enzyme complex. Crystal structures are extremely valuable as high-resolution snapshots of enzyme complex, but they lack the dynamic information about the enzyme complex that is required for a complete understanding of the autoactivation process. In addition, there is not crystal structure available of the [Aurora B/IN-box] complex in the unphosphorylated state and therefore, there is an important gap in understanding the conformational changes taking place during the activation process.

Proteins are dynamic macromolecules that exist in multiple conformations in their native state. An organized protein will explore a smaller number of conformations with less structural variability, than a disordered protein that will explore a larger number of conformations in the same time. In addition, there is an increasing evidence that the dynamic properties of an enzyme play a critical role in its catalytic functions (Ahuja et al., 2019; Tzeng and Kalodimos, 2012). For example, an increase in dynamics or local flexibility may facilitate the adoption of the different conformations required during the catalytic cycle (Boehr et al., 2006; Tzeng and Kalodimos, 2012; Xie et al., 2020). On the other hand, the transition from a highly dynamic, disordered enzyme to a more structured enzyme has also been documented as a form of regulation of enzyme catalysis (Wei et al., 2016; Xiao et al., 2015). In particular, an intensive line of research on protein kinases has shown that the internal dynamics associated with the movements of the enzyme’s subdomains regulate enzymatic activity (Ahuja et al., 2019; Pegram et al., 2021; Taylor and Kornev, 2011). While detailed dynamics analyzes are available for several eukaryotic protein kinases activated by phosphorylation (Foda et al., 2015; Wang et al., 2019), including the closely related Aurora A (Cyphers et al., 2017; Gilburt et al., 2017; Pitsawong et al., 2018; Ruff et al., 2018), the effect of phosphorylation on the dynamic properties of the [Aurora B/IN-box] complex is not yet known.

Here, we use a combination of experimental (hydrogen-deuterium exchange coupled to mass-spectrometry (HDX-MS) and kinetics) and computational (molecular dynamics; MD) approaches to follow the internal dynamics of the [Aurora B/IN-box] complex in solution -both in the unphosphorylated, inhibited, ([Aurora B/IN-box]^no-P^) and fully phosphorylated, active, ([Aurora B/IN-box]^all-P^) states. We found that in the absence of phosphorylation, there is structural disorder scattered along several regions of the enzyme complex. In contrast, in the fully phosphorylated enzyme complex, both Aurora B and the IN-box are structurally organized, resulting in a fully functional enzyme.

To investigate the contribution of individual phosphorylation sites to the dynamics and enzymatic activity, we used a protein ligation method to generate partially phosphorylated forms of the enzyme complex in which either only the activation loop of Aurora B ([Aurora B/IN-box]^loop-P^) or only the TSS motif of the IN-box ([Aurora B/IN-box]^IN-P^) is phosphorylated. Phosphorylation of each individual site in isolation has only a minor effect on enzymatic activity and structural stability, whereas simultaneous phosphorylation of the activation loop in Aurora B and the TSS-motif in the IN-box synergistically affects the structure and dynamics of the enzyme complex, leading to a dramatic increase in kinase activity. Further, we identify phosphorylation in the activation loop of Aurora B as a rate-limiting intramolecular event in autoactivation. Finally, only when both sites are phosphorylated are the global movements of the N- and C-lobes of the enzyme (opening and closing of the active site and twisting between the lobes) coordinated with the movements of the activation loop, resulting in a fully functional kinase.

Our combined experimental and theoretical approach has allowed us to study in parallel unphosphorylated, partially phosphorylated, and fully phosphorylated [Aurora B/IN-box] and to provide the missing information necessary to build a more detailed model of [Aurora B/IN-box] autoactivation. Our model is important for understanding the role of protein dynamics in achieving full enzyme activity, and it may also serve as a starting point for the development of new Aurora B-specific inhibitors.

## Results

### Autoactivation of [Aurora B/IN-box] by phosphorylation induces structural organization in multiple regions of the enzyme complex

The [Aurora B/IN-box] complex is activated by autophosphorylation, resulting in an enzyme with at least 100-fold higher enzymatic activity (Zaytsev et al., 2016) (fig. S1). To obtain information on the possible structural and dynamic changes underlying the observed differences in enzymatic activity between unphosphorylated [Aurora B/IN-box]^no-P^ and the fully phosphorylated enzyme complex, [Aurora B/IN-box]^all-P^, we monitored the exchange of hydrogen for deuterium (HDX) in the protein backbone of [Aurora B/IN-box] from *Xenopus laevis*. HDX is based on the fact that in aqueous solution, the amide protons of peptide bonds are exchanged with protons from the solvent (Mayne, 2016). Thus, by changing the solvent from H_2_O to D_2_O, HDX can be measured. The unstructured protein regions are the fast exchangers, while well-ordered regions where the amide backbone hydrogen atoms are involved in hydrogen bonds are exchanged more slowly. HDX analysis of proteins has proven useful in determining the activation mechanisms of numerous enzymes (Dawicki-McKenna et al., 2015; Habibi et al., 2019; Lorenzen and Pawson, 2014; Sours and Ahn, 2010; Zhang et al., 2020). By comparing the differences in deuterium uptake in the phosphorylated and unphosphorylated forms of the enzyme complex, we identified the regions of the enzyme where dynamic changes occur during phosphorylation. HDX measurements were performed using mass spectrometry (see Materials and Methods and fig. S2 for details).

We measured HDX in [Aurora B/IN-box]^no-P^ and [Aurora B/IN-box]^all-P^ at four time points. Comparison of HDX measurements for the two phosphorylation states reveals several regions of the enzyme complex with profound HDX differences (Fig. 1). The unphosphorylated form of the enzyme takes up more deuterium in all regions where HDX differences were detected, indicating greater flexibility and lower structural organization. Consequently, the fully active form, [Aurora B/IN-box]^all-P^, has a lower HDX, indicating a more rigid structural organization. A detailed comparison between the two enzyme forms reveals four regions with significant HDX differences (see Materials and Methods) in Aurora B (indicated by orange circles 1 to 4 in Fig. 1A) and one region in the IN-box (indicated by the green circle in Fig. 1A).

**Figure 1.**
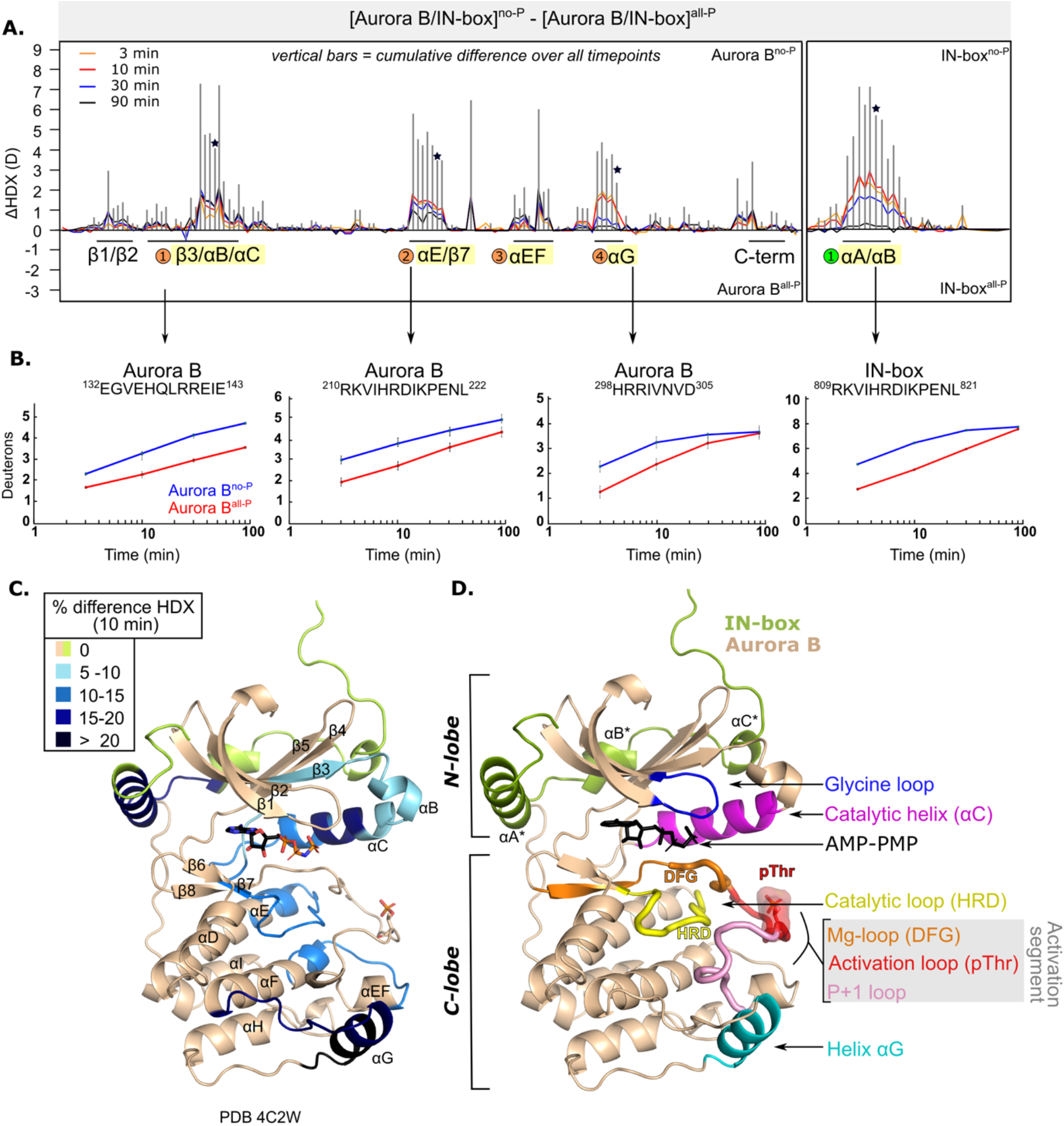
After phosphorylation, the [Aurora B/IN-box] complex becomes more rigid. **A**. Butterfly-difference plot for a comparison of H/D exchange differences between [Aurora B/IN-box]^no-P^ and [Aurora B/IN-box]^all-P^, measured over 4 time points (orange – 3 min; red - 10 min; blue - 30 min; black - 90 min). Each of the vertical bars represents a single peptide arranged according to its position from the N- to the C-terminus. The size of the vertical bars represents the cumulative difference ([Aurora B/IN-box]^all-P^ - [Aurora B/IN-box]^no-P^) or ΔHDX over all time points. It is noticeable that the difference in deuterium uptake is mainly positive, indicating an overall faster exchange and more flexible structure in [Aurora B/IN-box]^no-P^ than in [Aurora B/IN-box]^all-P^ for both Aurora B and IN-box. Secondary structural elements that are encompassed by peptides with differences in deuterium uptake are indicated below the baseline. Structural elements containing peptides with significant ΔHDX are highlighted in yellow and indicated with orange circles for Aurora B and green circles for the IN box. The asterisk indicates representative peptides for which uptake plots are shown in B.**B**. The uptake plots for representative peptides with high uptake differences. **C**. HDX differences between [Aurora B/IN-box]^no-P^ and [Aurora B/IN-box]^all-P^ captured after 10 minutes of exchange and mapped on ribbon diagram of the [Aurora B/IN-box] complex (PDB 4C2W). The Aurora B secondary structure elements are labeled. **D**. Ribbon diagram of [Aurora B/IN-box] with Aurora B in beige and IN-box in green. The secondary structure elements of the IN-box (αA*, αB* and αC*) are labeled. The important catalytic and regulatory elements of Aurora B are highlighted: glycine loop (blue), catalytic helix αC (magenta), catalytic loop with histidine-arginine-aspartic acid (HRD) motif (yellow) and helix αG (cyan). The active segment consists of an Mg-loop with DFG motif (aspartic acid-phenylalanine-glycine) (orange), an activation loop with phosphorylated threonine (red), and a P+1 loop (pink) involved in peptide substrate binding.

In Aurora B, the first region spans residues 120 to 156 and includes part of the Δ3-strand, the αB-helix, the catalytic αC-helix, and the loop region following the αC-helix (Fig. 1 A-C). This portion of Aurora B includes the residues Aurora B^Lys122^ and Aurora B^Glu141^, which play a role in defining the active conformation of the enzyme (Sessa et al., 2005). The middle portion of the αC helix is particularly rigid in the phosphorylated enzyme complex (Fig. 1C) and shows > 15% HDX protection across all time points. It is known from studies of other eukaryotic protein kinases that the αC-helix plays a critical role in the organization of the productive enzyme complex (Kornev and Taylor, 2015). This helix contributes to the positioning of the activation segment, which includes the magnesium-binding loop (DFG loop), the activation loop with phosphorylated Aurora B^Thr248^, and the P+1 loop (Fig. 1D). The αC-helix residues also contribute to the formation of the R-spine, a series of hydrophobic interactions extending from the N-lobe to the C-lobe of protein kinases, that are critical for the adoption of an active conformation (Kornev and Taylor, 2015). In addition, helices αB and αC provide an interacting surface for binding regulators of kinase activity in a variety of protein kinases (Leroux and Biondi, 2020) and are a site of interaction between the structurally similar Aurora A kinase and its binding partner TPX2 (Bayliss et al., 2003).

The second region spans residues 205 to 223 and includes the last five amino acids of the αE-helix, a linker region that includes the major catalytic loop (HRD loop) and the β7-strand. The third region spans residues 254 to 272 and includes the end of the activation segment, the αEF-helix, the loop upstream of the αF-helix, and the first turn of the αF-helix, which is an important organizer of the C-lobe (Kornev et al., 2008). The fourth region spans residues 284 to 305 and includes the αG-helix and the preceding loop. This region is part of the peptide substrate recognition site and is thought to play a regulatory role in protein kinases (Ahuja et al., 2019).

The IN-box is highly dynamic with a very fast HDX rate in both [Aurora B/IN-box]^no-P^ and [Aurora B/IN-box]^all-P^ (most of the peptides are completely exchanged at the early time points, Data set D1). The only segment of IN-box where differences in deuteration are detected extends from position 805 to 825 and contains the only well-defined secondary structure elements: IN-box^αA^ helix and the short IN-box^αB^ helix connected by a linker (Fig. 1A-D). HDX kinetics for this region shows a strong component of EX1 kinetics and a smaller component of common EX2 kinetics. The EX1 kinetics is reflected in a bimodal distribution with the clear appearance of two distinct isotopic envelopes of the masses (fig. S3). EX1 kinetics is observed in structural elements that undergo transitions between folded and unfolded states at a relatively slow refolding rate, resulting in HDX occurring in an all-or-nothing manner (Mayne, 2016). Although EX1 features are present in both, the unphosphorylated and phosphorylated, states of the enzyme complex, they are significantly faster and more pronounced in [Aurora B/IN-box]^no-P^ (fig. S3).

A possible explanation for the large deuterium uptake in the unphosphorylated [Aurora B/IN-box] could be the dissociation of the IN-box from Aurora B at the low μM concentration (0.4 μM) used in our HDX experiment. In this scenario, the interacting surfaces of Aurora B and IN-box would be available for H/D exchange. To determine whether the [Aurora B/IN-box] complex remained intact in our HDX experimental conditions, we used mass photometry to measure the molecular weight (MW) of the enzymatic complex, [Aurora B/IN-box]^no-P^ and [Aurora B/IN-box]^all-P^ at low nM concentrations. In both samples, we observed a single uniform peak corresponding to MW of ∼44 kD (MW_Aurora B_=35.4 kD; MW_IN-box_=8 kD), the measurements were performed at 10 nM enzyme concentration. This clearly confirms that the interactions between Aurora B and IN-box are stable under the HDX experimental conditions in the two states, phosphorylated and unphosphorylated, (fig. S4) and that the differences observed by HDX are due to phosphorylation-induced changes in enzyme dynamics rather than dissociation between the two polypeptide chains.

The higher degree of HDX protection detected in the active site and key regulatory regions of the enzyme (Fig. 1C and D) suggests that the structural organization that occurs in the phosphorylated enzyme, [Aurora B/IN-box]^all-P^, is important for enzymatic activity. Our results are consistent with similar studies on other protein kinases where phosphorylation of the threonine residue in the activation loop leads to rigidification of the structure (Lorenzen and Pawson, 2014; Steichen et al., 2010). To gain more detailed structural insight into the dynamics and structure of the [Aurora B/IN-box] complex and to better understand the specific role of the IN-box in regulating Aurora B kinase activity, we performed MD simulations.

### Molecular dynamics simulations reveal major conformational change in the IN-box upon enzyme complex phosphorylation

The HDX experiments show that phosphorylation has a clear effect on the dynamics of [Aurora B/IN-box], but do not provide information on the nature of the conformational and dynamic changes that take place. To gain insight into the structural rearrangements during the autoactivation process at atomic resolution, we performed a 2.8 μs molecular dynamics simulation (MD) of the enzyme complex in the phosphorylated, [Aurora B/IN-box]^all-P^, and the unphosphorylated, [Aurora B/IN-box]^no-P^, using the available X-ray structure (PDB 4C2W) (Sessa and Villa, 2014) as a template (see Materials and Methods and Fig.2). This is a crystal structure of the *X. laevis* [Aurora B/IN-box] complex in which the activation loop is phosphorylated, and the IN-box contains residues 797-847 (ends just before the TSS motif). The structure contains an ATP analog (ANP-PMP) in the active site (instead of bulky inhibitors) and represents an enzyme of the same species as our experimental system, so we consider it the most appropriate starting point for our MS simulations. The C-terminus of the IN-box was extended by the residues IN-box^848-856^ (dashed line in the Fig. 2A), to correspond exactly to our experimental enzyme complex, and the AMP-PMP in the active site was replaced by ATP -Mg. To obtain [Aurora B/IN-box]^no-P^, the phosphate group of Aurora B^Thr248^ was removed, whereas for [Aurora B/IN-box]^all-P^, phosphate groups at IN-box^Ser849^ and IN-box^Ser850^ were added (Fig. 2A; see Materials and Methods for details). During the simulation, the systems were stable and retained the tertiary and quaternary structures in the folded regions of the complex (fig. S5A). The major structural differences between the two forms of the enzyme complex were observed in the conformation of the C-terminal part of IN-box and in the conformation of the activation loop.

**Figure 2.**
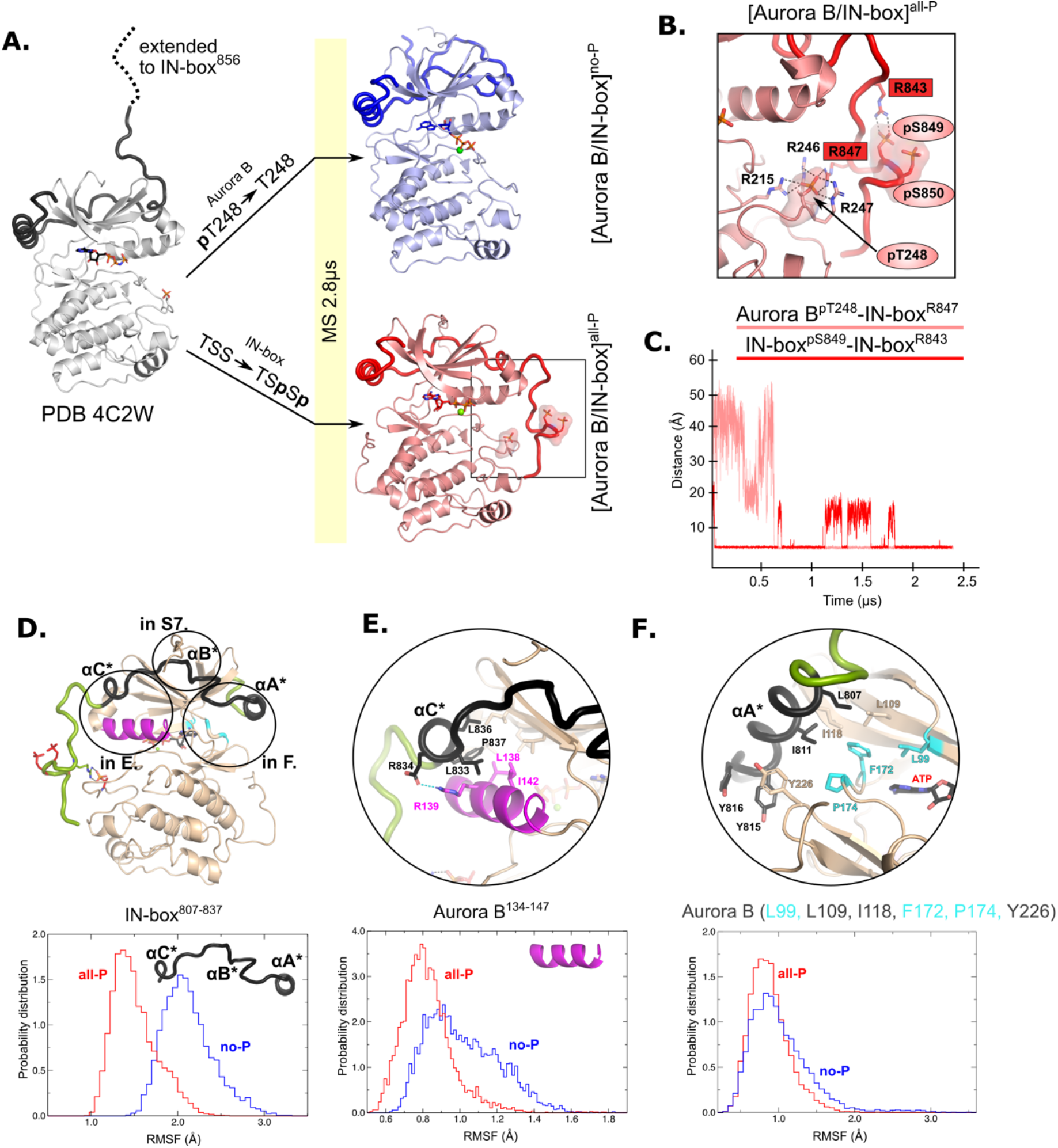
MD simulation of [Aurora B/IN-box]^no-P^ and [Aurora B/IN-box]^all-P^ reveals large conformational changes in the C-terminal part of IN-box and dynamic coupling between IN-box and the Aurora B catalytic helix. **A**. Initial models for MD stimulation were generated from PDB 4C2W (ribbon diagram in gray; IN-box is shown as a black coil). The C-terminal part of IN-box was extended to the end (to include up to IN-box^856^; dotted line), AppNHp was replaced by ATP-Mg and for [Aurora B/IN-box]^no-P^, the phosphate group of Aurora B^pT248^ was removed. For [Aurora B/IN-box]^all-P^, phosphates were added to IN-box^Ser849^ and IN-box^Ser850^. Simulations for both phosphorylation states were run for 2.8 μs and the ribbon diagram for the final structure for [Aurora B/IN-box]^no-P^ is shown in blue and the final structure for [Aurora B/IN-box]^all-P^ is shown in red (IN-box is shown as coil in a darker color). The interactions within the boxed region of the [Aurora B/IN-box]^all-P^ structure are shown in B. **B**. Enlarged view showing the interactions of Aurora B^pThr248^ with residues in Aurora B and IN-box in the final MD confirmation of [Aurora B/IN-box]^all-P^. **C**. The distance between the phosphorus atom in Aurora B^pThr248^ and the Cζ atom in IN-box^Arg847^ (pink) and between the phosphorus atom in IN-boxp^Ser847^ and the Cζ atom in IN-box^Arg843^ (red) during the MD simulation. The Cζ atom was chosen for the distance measurements because it is bonded to all three nitrogen atoms in the arginine residue, which can form a hydrogen bond with phosphate. **D**. The ribbon diagram of the [Aurora B/IN-box]^all-P^ structure (rotated 180° compared to the orientation in A). Aurora B is in beige and IN-box is in green. The N-terminal part of IN-box, IN-box^807-837^, which contains well-defined structural elements, αA*, αB*, αC*, is colored black. Parts of Aurora B that interact with this N-terminal IN-box and are important for catalysis are in magenta (Aurora B^αC^) and cyan (residues forming the ATP binding site). The interfaces between three parts of the IN-box are circled and shown in more detail in panels E, F and Supplementary figure S7. Below is the plot of probability density in the y axis versus the RMSF for IN-box^807-837^. RMSF in [Aurora B/IN-box]^all-P^ is in red and [Aurora B/IN-box]^no-P^ is in blue. **E**. Interface between IN-box^αC*^ and Aurora B. Below is the plot of probability density distribution vrs RMSF for Aurora B^αC^ with Same labeling as in D. **F**. Interface between IN-box^αA*^ and Aurora B. Three residues in Aurora B that form the ATP binding site are cyan. Below is the same type of plot as is D and E for for Aurora B (Leu99, Leu109, Ile118, Phe172, Pro174 and Tyr226).

The C-terminal region of the IN-box, which is not visible in the X-ray structure of Aurora B from *X. laevis* (Sessa and Villa, 2014) was first modeled as an unfolded coil in solution, pointing away from the enzyme body and not interacting with any part of the enzyme complex (Fig. 2A, dashed line). In the MD-simulation of [Aurora B/IN-box]^no-P^, where neither Aurora B^Thr248^ nor the TSS-motif of IN-box are phosphorylated, the IN-box showed great flexibility. Its C-terminal part remains unstructured and does not stably interact with Aurora B throughout the simulation period (Fig. 2A and movie M1).

On the contrary, the simulations of phosphorylated [Aurora B/IN-box] showed that the presence of phosphates in Aurora B and IN-box rapidly stabilizes a defined conformation in which the C-terminal region of IN-box interacts with the Aurora B activation loop (Fig. 2A and movie M2). At the beginning of the simulation, two phosphates of IN-box (IN-box^pSer849^ and IN-box^pSer850^) are immediately stabilized by adjacent positively charged residues, IN-box^Arg843^ and IN-box^Lys846^. Moreover, IN-box^Arg847^ forms an electrostatic interaction with the phospho-threonine of the activation loop, Aurora B^pThr248^, early in the simulation (0.7 μs), and this interaction remains stable until the end of the simulation (the next 1.8 μs) (Fig. 2B and C). In addition to IN-box^Arg847^, Aurora B^pThr248^ is stabilized by three arginine residues from Aurora B -two adjacent residues in the activation loop, Aurora B^Arg246^ and Aurora B^Arg247^, and Aurora B^Arg215^ from the catalytic loop (HRD). Thus, stabilization of negative phosphate in the [Aurora B/IN-box]^all-P^ activation loop involves interaction with four arginine residues and is more extensive than phosphate stabilization in the initial structure (PDB 4C2W), in which Aurora B^pThr248^ is present but the IN-box TSS motif is absent (fig. S5B). In the initial structure, Aurora B^pThr248^ is stabilized by only two Aurora B arginines, Aurora B^Arg246^ and Aurora B^Arg215^, while Aurora B^Arg247^ faces the solvent. Interestingly, in the crystal structure of the phosphorylated human [Aurora C/IN-box] containing the phosphorylated IN-box TSS motif, the arrangement of the Aurora B and IN-box residues around the phosphorylated Aurora B threonine is similar (fig. S5C). In both, our simulation and the crystal structure of phosphorylated [Aurora C/IN-box], the IN-box forms stable interactions with the activation loop of Aurora B, which could lead to reduced dynamics and a more compact enzyme.

### Experimental (HDX) and theoretical (MD) approaches imply that the dynamics of IN-box and Aurora B are interconnected

Both HDX and MD clearly show that the IN-box is highly disorganized in the unphosphorylated state of the enzyme complex. However, the large differences in the dynamics of the IN-box observed in MD simulations in the C-terminal part of the IN-box are not detectable in our HDX experiments. There could be two reasons for this apparent discrepancy. First, the C-terminal part of the IN-box is anchored to Aurora B only by side-chain interactions, whereas the main-chain hydrogens do not form stable hydrogen networks, leaving this region free for H/D exchange despite a different conformation (fig. S5D). An alternative explanation is that HDX differences exist in the C-terminal region of the IN-box between two states, but the kinetics of the exchange occur in the millisecond range, which cannot be measured with our experimental setup.

Further, the N-terminal part, which has well-defined secondary structure elements, shows partial EX1 behavior, indicating partial unfolding (fig. S3). The N-terminal segment of the IN-box, which surrounds the N-lobe of Aurora B with multiple interactions, is the only part of the IN-box that is always associated with Aurora B. We wondered whether the dynamics of the interacting neighboring regions in Aurora B are coupled to the dynamics of the IN-box in the MD simulation.

We calculated the root mean square fluctuations (RMSF) during MS simulations for the N-terminal part of the IN-box, IN-box^807-837^, in the two forms of the enzyme complex (black coil in Fig. 2D). Higher RMSF are associated with larger structural deformations. Consistent with the HDX results, the RMSFs for this region are significantly higher in [Aurora B/IN-box]^no-P^ than in [Aurora B/IN-box]^all-P^ (Fig. 2D, graph). Next, we analyzed the RMSFs for the Aurora B regions interacting with the N-terminal fragment of the IN-box (Fig. 2E and F, fig S6). These regions include Aurora-B kinase elements essential for catalysis and substrate binding: the Aurora B^αC^, catalytic helix (Fig. 2E, magenta), and the region forming the ATP-binding site (Fig. 2F, cyan), respectively. In the dephosphorylated state, Aurora B^αC^ exhibits a very broad RMSF distribution shifted toward higher values, reflecting strong helix deformations. In contrast, the RMSF values in [Aurora B/IN-box]^all-P^ show a narrow distribution, clearly indicating a more structured Aurora B^αC^. Similarly, the region of Aurora B that is forming the ATP binding site and is in contact with IN-box^αA^ shows a broader distribution of RMSF shifted to higher values, in the dephosphorylated state (Fig. 2F). In summary, the Aurora B regions in contact with the N-terminal IN-box segment are structurally more disordered in the absence of phosphorylation than when the enzyme complex is fully phosphorylated, strongly suggesting that the dynamics of the adjacent regions of Aurora B and IN-box are coupled. In the unphosphorylated form of the enzyme complex the structural disorder spreads from one partner to the other. Therefore, the IN-box could act as a negative regulator of the kinase activity by transferring disorder to the kinase domain. Conversely, structural order is imposed on both Aurora B and the IN-box by phosphorylation.

However, the [Aurora B/IN-box]^all-P^ complex is phosphorylated in both Aurora B (activation loop – Aurora B^Thr248^) and the IN-box (TSS motif – IN-box^pSer849^ and IN-box^pSer850^), so it is not clear whether the observed changes are the result of phosphorylation of the Aurora B activation loop or the TSS motif in the IN-box, or both. We next wanted to determine the contribution of each of these phosphorylation sites to the dynamics of the enzyme complex and to its enzymatic activity.

### Generation of partially phosphorylated [Aurora B/IN-box] complexes

Previous attempts to uncouple the effects of the two phosphorylation sites in the [Aurora B/IN-box] enzyme complex were based on the analysis of phospho-null mutants (Abdul Azeez et al., 2019; Wang et al., 2011) in which the TSS motif was changed to TAA. Although the mutational approach to mimic a phospho-null IN-box was possible, phospho-mimetic mutations in which the threonine in the activation loop or the serine residues in the IN-box TSS-motif are replaced by glutamic acids to mimic phosphorylation resulted in catalytically dead enzymes, contrary to expectations. Phospho-null mutations in the activation loop also resulted in a dead kinase. Since it was not possible to obtain enzyme complexes with phosphorylation only in the IN-box, the contribution of phosphorylation of the activation loop to enzyme activity could not be understood.

Because the IN-box is relatively close to the C-terminal end of the construct this region of the protein is suitable for protein ligation. Therefore, we used a synthetic peptide approach to introduce posttranslational modifications (Ghosh et al., 2011) (Fig. 3A). We generated a truncated IN-box^790-844^ (IN-box^ΔC^) fused to an intein that was co-expressed with Aurora B in a similar manner to the original [Aurora B/IN-box] construct. After purification of the complex and intein cleavage, the resulting [Aurora B/IN-box]^IN-ΔC^ was ligated with a synthetic peptide containing the IN-box residues IN-box^845-858^ including the TSS motif (Fig 3B). The resulting protein differs from the original construct by a single amino acid that remains as a “scar” of the ligation – IN-box^Phe845^ becomes IN-box^Cys845^ (indicated by the asterisk in Fig. 3A). The enzyme complex containing phosphorylated Aurora B^Thr248^ in the activation loop and the unphosphorylated IN-box, [Aurora B/IN-box]^loop-P^, was obtained by ligation of the [Aurora B/IN-box]^IN-ΔC^ with a peptide containing the unphosphorylated IN-box^845-858^ (Fig. 3B). To ensure that Aurora B^Thr248^ was phosphorylated, the [Aurora B/IN-box]^IN-ΔC^-intein was incubated with traces of the wild-type [Aurora B/IN-box]^all-P^ complex (WT) before ligation. The WT enzyme was removed in a purification step before ligation (see Materials and Methods for further details). To generate the enzyme complex phosphorylated only in the TSS motif of the IN-box, [Aurora B/IN-box]^IN-P^, the truncated [Aurora B/IN-box]^IN-ΔC^-intein complex, was incubated with λ-phosphatase to ensure that Aurora B^Thr248^ was not phosphorylated. After further purification to remove the λ-phosphatase, the unphosphorylated [Aurora B/IN-box]^IN-ΔC^ was ligated with an IN-box^845-858^ phosphorylated at the two serine residues of the TSS motif. In all cases, size exclusion chromatography was performed as a final purification step to ensure that unligated peptides were removed (fig. S7), whereas the presence or absence of phosphates in the Aurora B^Thr248-P^ or TSS motif in the IN-box was confirmed by mass spectrometry (fig. S8).

**Figure 3.**
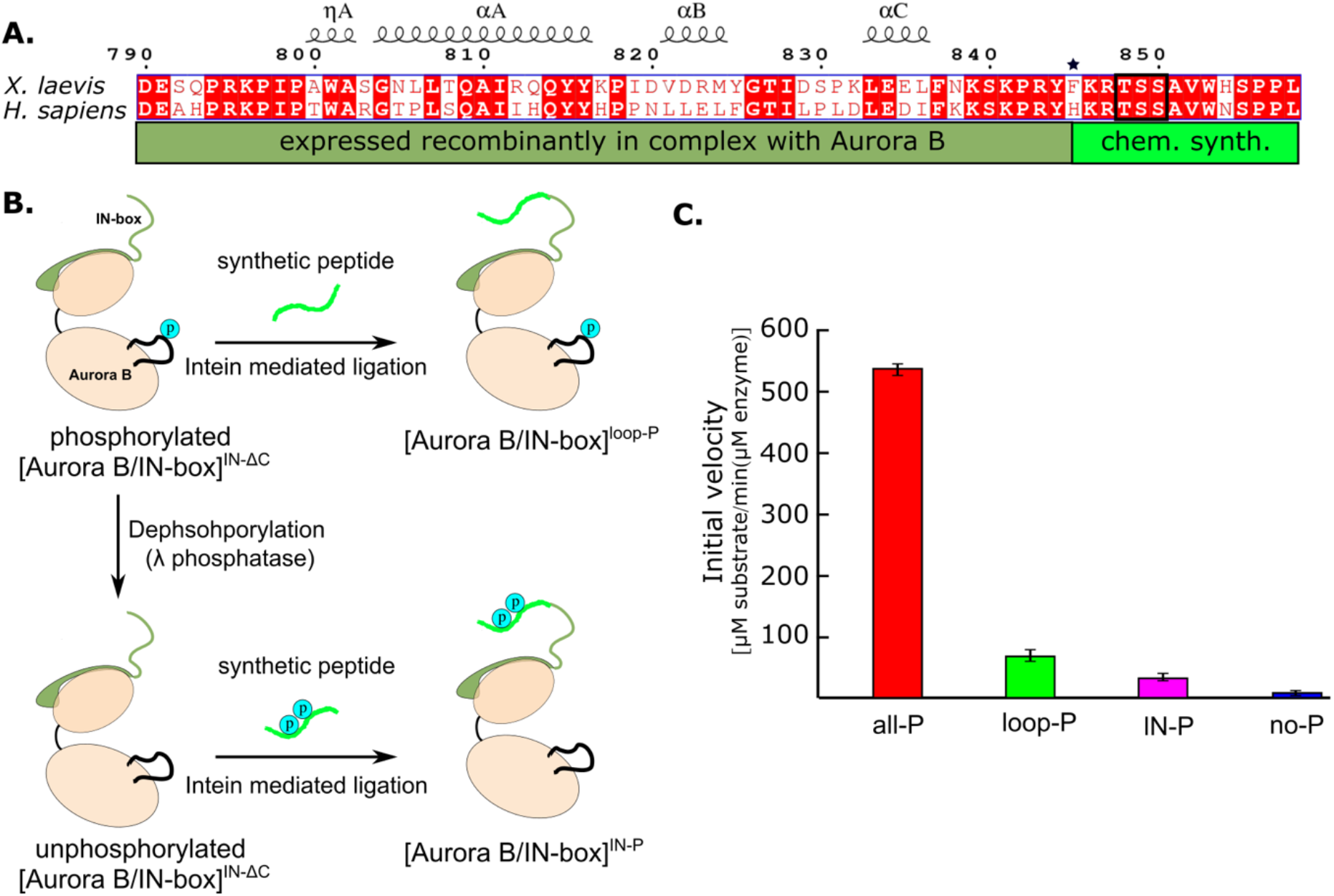
Design and kinetic analysis of the partially phosphorylated [Aurora B/IN-box] complex. **A**. Sequence overlay of the IN-box, region of INCENP, from *H. sapiens* and *X. laevis*. The conserved amino acids are shown in red and the elements of the secondary structure are at the top. Residues 790-844 (dark green) are expressed together with Aurora B, in a bicistronic construct, giving rise to [Aurora B/IN-box]^IN-ΔC^, which is then ligated to a synthetic peptide containing residues 846-858 (light green). The synthetic peptide contains a TSS motif that can be phosphorylated (black box). The Aurora B^Phe845^ (indicated by asterisks) is converted to Cys during ligation. **B**. The schematic shows the steps to prepare [Aurora B/IN-box]^loop-P^ and [Aurora B/IN-box]^IN-P^ using [Aurora B/IN-box]^IN-ΔC^ and synthetic peptides. **C**. Enzymatic activity determined from initial velocities for [Aurora B/IN-box]^all-P^ (red), [Aurora B/IN-box]^loop-P^ (green), [Aurora B/IN-box]^IN-P^(magenta) and [Aurora B/IN-box]^no-P^ (blue).

Having prepared semisynthetic, partially phosphorylated intermediates, we wanted to test whether they could reach their full enzymatic activity when autophosphorylated in the presence of ATP-Mg. Indeed, both semisynthetic constructs showed similar initial rates of enzymatic activity as the wild type [Aurora B/IN-box]^all-P^ after incubation with ATP-Mg (fig. S9A). This gave us confidence that neither the single mutation nor the process of chemical ligation affected the intrinsic nature of the enzyme complex.

### The two phosphorylation sites synergistically contribute to Aurora B kinase activity

Next, we compared the initial rates of enzymatic reaction for the partially phosphorylated enzyme complexes with fully phosphorylated, [Aurora B/IN-box]^all-P^, and unphosphorylated, [Aurora B/IN-box]^no-P^, in the presence of high substrate peptide concentrations (600 μM) (Fig. 3C). Interestingly, although both partially phosphorylated enzyme complexes showed detectable enzymatic activity, their initial rates were much lower than those of the fully active enzyme, [Aurora B/IN-box]^all-P^. [Aurora B/IN-box]^loop-P^ exhibits only ∼10 % and [Aurora B/IN-box]^IN-P^ only ∼5 % of the maximum initial rate. Although these values are still higher than the 1 % activity measured for the unphosphorylated enzyme, [Aurora B/IN-box]^no-P^, they are drastically lower than the maximum initial rates measured for the fully phosphorylated [Aurora B/IN-box]^all-P^. Thus, the simple addition of the activities of [Aurora B/IN-box]^loop-P^ and [Aurora B/IN-box]^IN-P^ accounts for only ∼15 % of the full enzyme activity, clearly showing that the phosphorylation of the Aurora B activation loop and the TSS motif of the IN-box act synergistically, rather than independently, to achieve the full activity of the [Aurora B/IN-box] complex.

For the [Aurora B/IN-box]^loop-P^ construct, we determined the kinetic constants, v_max_ and K_M_ for the peptide substrate (fig. S9B). The v_max_ of this enzyme complex is about threefold lower than that of the fully phosphorylated, [Aurora B/IN-box]^all-P^, complex, whereas K_M_ increases dramatically (10-fold) (Table 1). This result suggests that the absence of phosphates in the IN-box TSS motif has a strong effect on substrate binding and a moderate effect on catalysis of the reaction and supports the role of the IN-box in stabilizing the open conformation of the activation segment that forms the peptide-substrate binding site. Overall, absence of phosphates in IN-box results in a twenty-fold decrease in the catalytic efficiency (k_cat_/K_M_) of [Aurora B/IN-box]^loop-P^ compared to the fully phosphorylated complex. Similar kinetic behavior was observed by Azeez and coworkers (Abdul Azeez et al., 2019) in a construct where TSS was replaced with TAA.

**Table 1.**
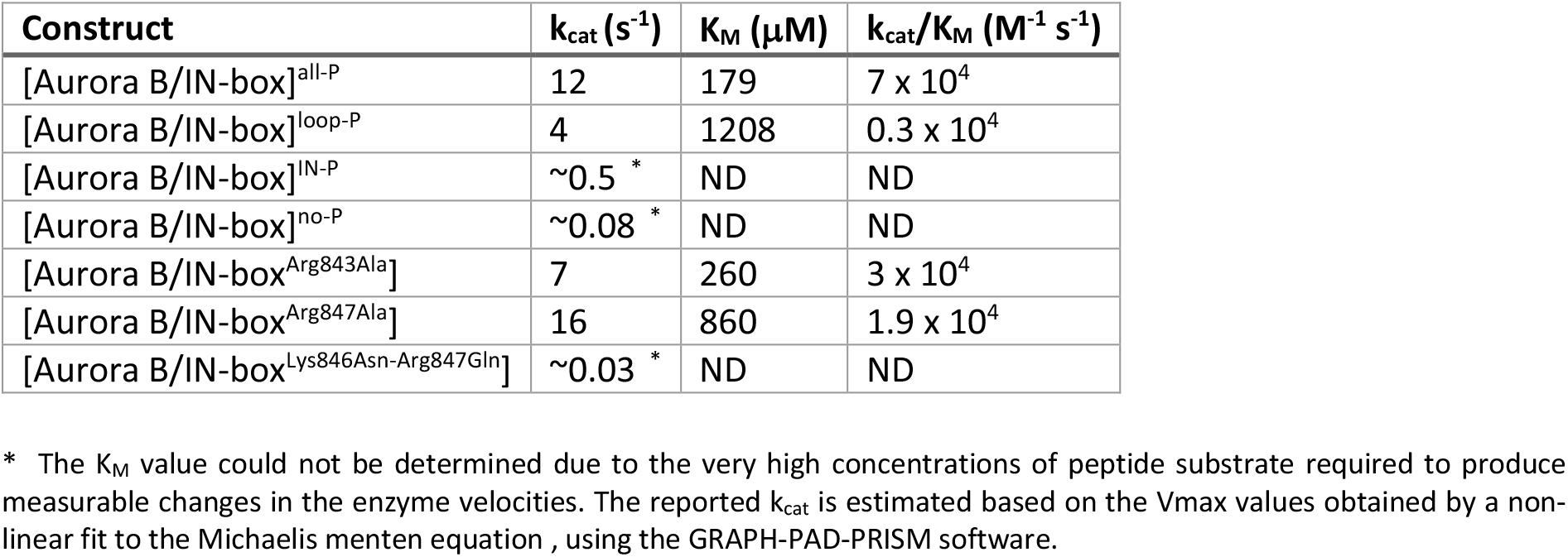
Kinetic constants for [Aurora B/IN-box] complexes.

| Construct | $k_{\text{cat}}$ ($\text{s}^{-1}$) | $K_M$ ($\mu\text{M}$) | $k_{\text{cat}}/K_M$ ($\text{M}^{-1} \text{s}^{-1}$) |
| --- | --- | --- | --- |
| [Aurora B/IN-box]<sup>all-P</sup> | 12 | 179 | $7 \times 10^4$ |
| [Aurora B/IN-box]<sup>loop-P</sup> | 4 | 1208 | $0.3 \times 10^4$ |
| [Aurora B/IN-box]<sup>IN-P</sup> | $\sim 0.5$ * | ND | ND |
| [Aurora B/IN-box]<sup>no-P</sup> | $\sim 0.08$ * | ND | ND |
| [Aurora B/IN-box<sup>Arg843Ala</sup>] | 7 | 260 | $3 \times 10^4$ |
| [Aurora B/IN-box<sup>Arg847Ala</sup>] | 16 | 860 | $1.9 \times 10^4$ |
| [Aurora B/IN-box<sup>Lys846Asn-Arg847Gln</sup>] | $\sim 0.03$ * | ND | ND |

\* The $K_M$ value could not be determined due to the very high concentrations of peptide substrate required to produce measurable changes in the enzyme velocities. The reported $k_{\text{cat}}$ is estimated based on the $V_{\text{max}}$ values obtained by a non-linear fit to the Michaelis menten equation , using the GRAPH-PAD-PRISM software.

We attempted to determine kinetic constants for [Aurora B/IN-box]^IN-P^, but this was not possible because a saturating concentration of the substrate peptide could not be obtained for this partially phosphorylated enzyme complex.

In summary, the experiments with the phosphorylated intermediates allowed us to unambiguously determine the contribution of each phosphorylation site independently and conclude that the phosphorylations of the Aurora B activation loop and the IN-box sites contribute synergistically to the kinase activity.

### Phosphorylation of each subunit of the complex leads to partial rigidification of the entire complex, but only the fully phosphorylated enzyme complex achieves complete structuring

We wanted to investigate whether the impaired catalytic activity in the partially phosphorylated enzyme complexes correlates with their structural organization, and therefore measured HDX in the [Aurora B/IN-box]^loop-P^ and the [Aurora B/IN-box]^IN-P^ enzyme complexes.

Because our partially phosphorylated complexes have the IN-box^Phe845Cys^ mutation, we first asked whether [Aurora B/IN-box^Phe845Cys^] exhibits the same HDX trends as [Aurora B/IN-box] when unphosphorylated and fully phosphorylated. Indeed, [Aurora B/IN-box^Phe845Cys^] shows higher dynamics in the unphosphorylated state than in the phosphorylated state with similar pattern as the enzyme complex without the mutation (fig. S10). We also observe the EX1 behavior in the N-terminal part of IN-box (fig. S11). In order to properly account for the contribution of the F845C mutation to the HDX differences, the reported values for [Aurora B/IN-box]^all-P^ and [Aurora B/IN-box]^no-P^ are measured on the enzyme complex carrying a mutation. The use of the mutant, [Aurora B/IN-box^Phe845Cys^], in the HDX experiments is labeled with a star, [Aurora B/IN-box]^all-P*^ and [Aurora B/IN-box]^no-P*^.

Both partially phosphorylated enzyme forms exhibited intermediate levels of HDX protection (Fig. 4 and figs. S12-13). Each of the phosphorylations alone results in intermediate structuring of the entire enzyme complex compared with the fully phosphorylated enzyme. For example, two Aurora B regions defining the catalytic site, the catalytic helix (Aurora B^αC^) and the catalytic loop (HRD loop), each exhibit partial H/D protection in both partially phosphorylated forms of the enzyme complex, corresponding to approximately half of the full protection observed in the fully phosphorylated active enzyme complex (Fig. 4B, first two peptides). The only exceptions to this rule are two regions that show equal or even greater H/D protection upon partial phosphorylation (black boxes in Fig. 4A and Fig. 4B, last two peptides). In the [Aurora B/IN-box]^IN-P^ complex, only the IN-box^αA^ helix shows high HDX protection. Interestingly, rigidification of the the IN-box^αA^ only partially affects neighboring Aurora B parts (see Fig.4A and 4B, first peptide encompassing Aurora B^αC^) suggesting that structuring of the IN-box is not sufficient to cause complete rigidification of the neighboring Aurora B regions. In the [Aurora B/IN-box]^loop-P^ complex, only the Aurora B^αG^-helix is strongly protected in HDX experiments (Fig. 4B). We further confirmed that this rigidification is independent of phosphorylation of the TSS motif in the IN-box, by measuring HDX in the enzyme complex in which TSS is mutated to TAA (fig. S14).

**Figure 4.**
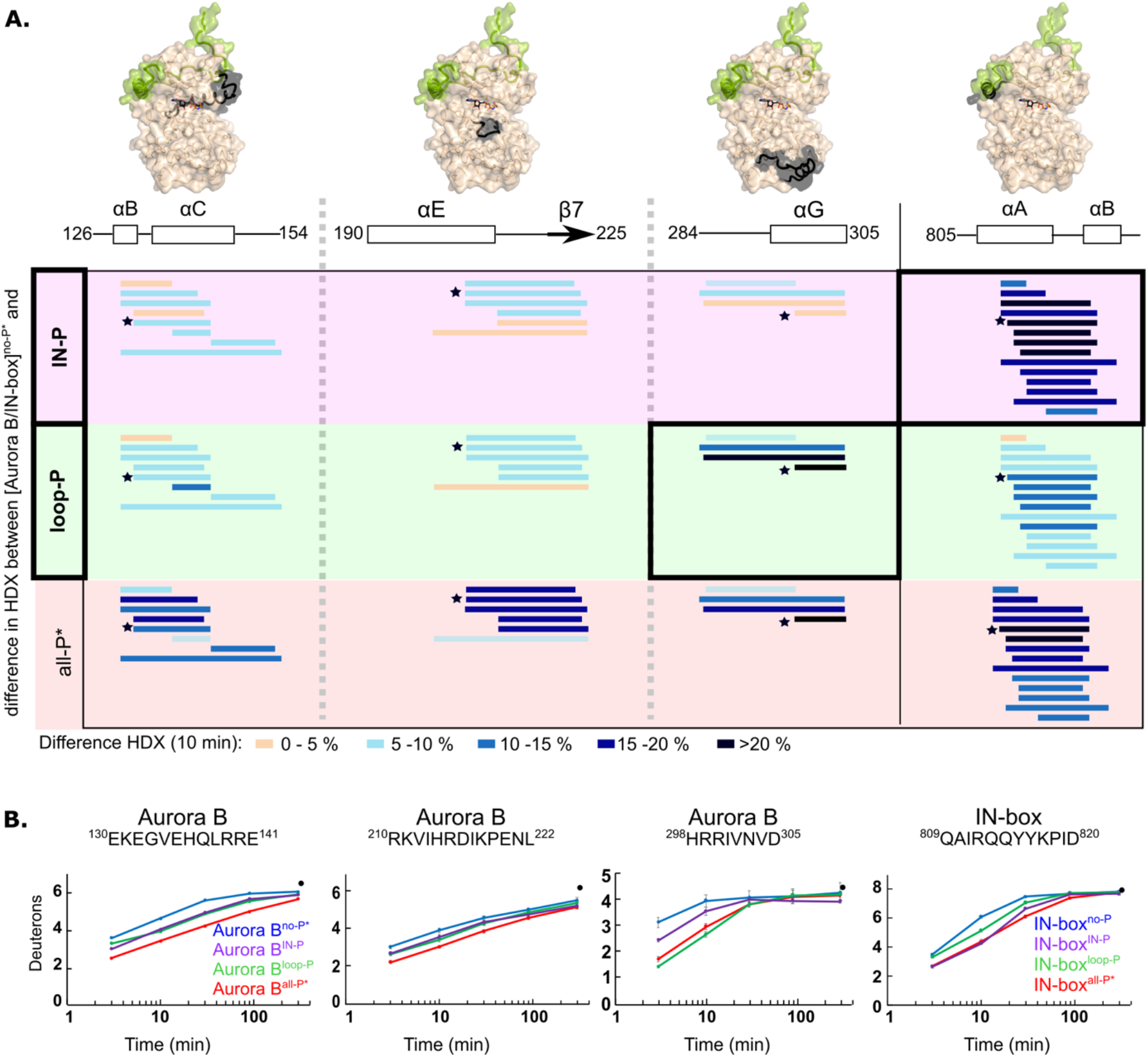
Two phosphorylations are required for maximal HDX protection in [Aurora B/IN-box] with only two exceptions in Aurora B^αG^ and IN-box^αA-αB^. **A**. Surface representation of the [Aurora B/IN-box] molecule (PDB 4C2W), with regions where HDX differences were detected shown in black. A difference plot of HDX uptake between [Aurora B/IN-box]^no-P*^ and either [Aurora B/IN-box]^loop-P^ (first row) or [Aurora B/IN-box]^IN-P^ (second row) after 10 minutes is shown for peptides in three different regions of Aurora B and one region in IN-box (these regions show the largest differences and are the same as in Fig. 1). For reference, the differences in HDX uptake between [Aurora B/IN-box]^no-P*^ and [Aurora B/IN-box]^all-P*^ are shown in the third row (they both carry the same “scar” mutation as partially phosphorylated intermediates to calibrate for the effects of the mutation on enzyme dynamics). Peptides are shown as horizontal lines and colored according to differences in deuterium uptake (see legend below). The asterisks indicate representative peptides for which uptake plots are shown in B. Note that in [Aurora B/IN-box]^loop-P^ only Aurora B^αG^ shows HDX protection comparable to [Aurora B/IN-box]^all-P^, while in [Aurora B/IN-box]^IN-P^ this is only true for IN-box^αA^. **B**. The uptake plots for representative peptides from A. The black dot is showing HDX for fully deuterated peptide under same experimental condition.

Thus, HDX analysis shows that phosphorylation in the activation loop and phosphorylation in the IN-box each individually affect the entire enzyme complex, but full structural stability is achieved only when both units are phosphorylated, consistent with the synergistic interaction inferred from enzyme kinetics experiments.

### Phosphorylation of the Aurora B activation loop triggers a conformational change in the C-terminal region of the IN-box

To understand how partial phosphorylation affects enzyme structure, we next performed MD simulations for partially phosphorylated enzyme complexes in the same manner as for unphosphorylated and fully phosphorylated [Aurora B/IN-box] (Fig. 5A and fig. S15A).

**Figure 5.**
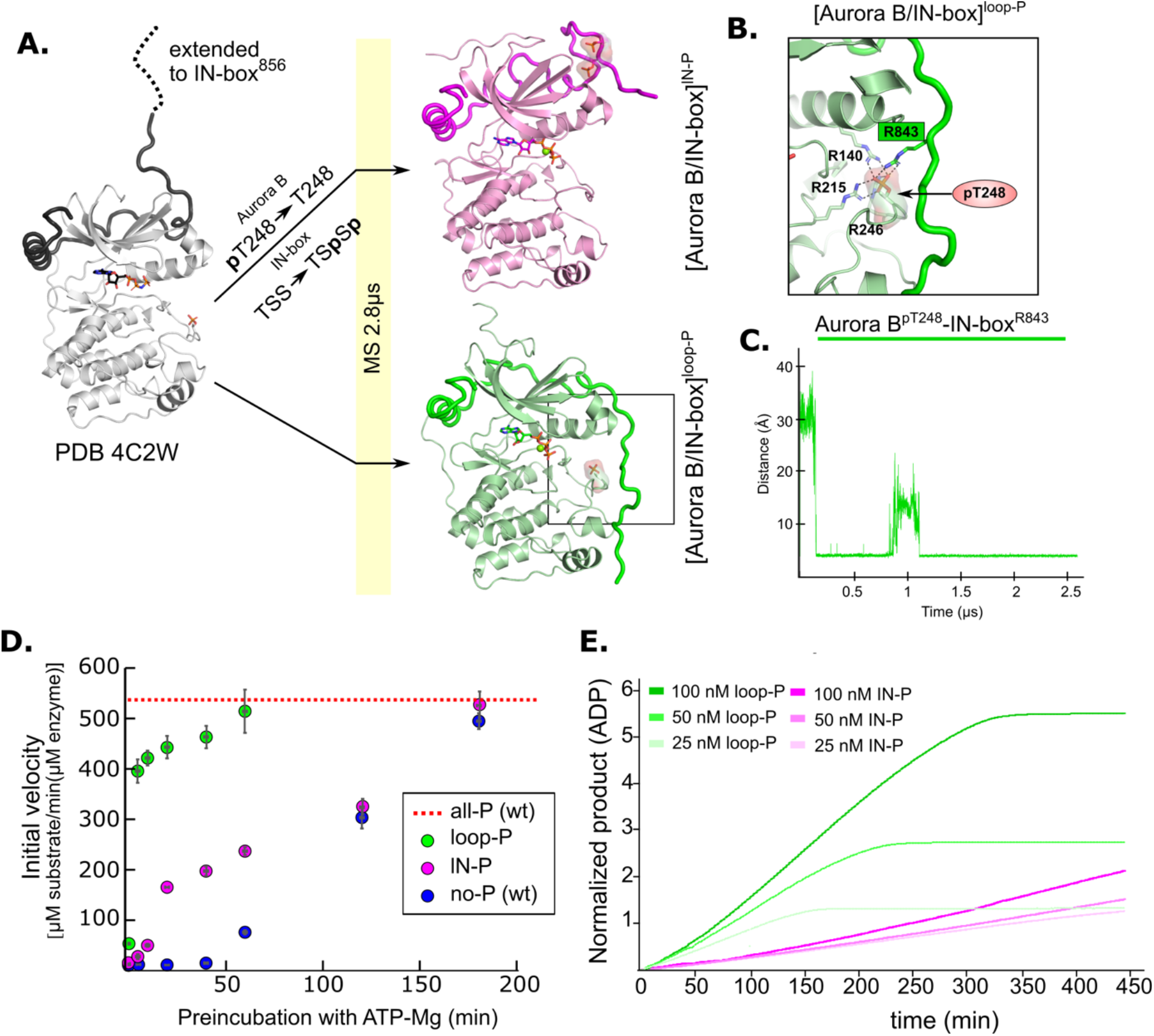
MD simulation and kinetic experiments show that phosphorylation of the Aurora B activation loop triggers a conformational change in the C-terminal region of the IN-box priming the [Aurora B/IN-box] complex for the activation. **A**. Initial models for MD stimulation were generated as previously described (see legend to Fig. 1). For [Aurora B/IN-box]^IN-P^, the phosphate group of Aurora B^pT248^ was removed and phosphates were added to IN-box^Ser849^ and IN-box^Ser850^. For [Aurora B/IN-box]^loop-P^, no phosphates needed to be added or removed because the initial structure already contained Aurora B^pThr248^. Simulations were run for 2.8 μs and ribbon diagram of the final structure for [Aurora B/IN-box]^IN-P^ is shown in magenta and the final structure for [Aurora B/IN-box]^loop-P^ is shown in green (IN-box is shown as a coil in a darker color). The interactions, within the boxed region of the [Aurora B/IN-box]^loop-P^ structure, are shown in B. **B**. Enlarged view showing the interactions of Aurora B^pThr248^ with residues in Aurora B and IN-box in the final MD confirmation of [Aurora B/IN-box]^loop-P^. **C**. The change in the distance between the phosphorus atom in Aurora B^pThr248^ and the Cζ atom in IN-box^Arg843^ during the MD simulation. **D**. Autoactivation assay for unphosphorylated or partially phosphorylated complexes, The enzyme was first pre-incubated with ATP-Mg at different times, followed by an enzymatic assay to determine the initial rate of substrate peptide phosphorylation. The plot shows the initial rates of enzymatic activity as a function of incubation time with ATP for [Aurora B/IN-box]^no-P^ (blue), [Aurora B/IN-box]^loop-P^ (green) and [Aurora B/IN-box]^IN-P^ (magenta). The full enzymatic activity of [Aurora B/IN-box]^all-P^ is shown as a red dotted line. Initial velocity measurements after different ATP preincubation times were performed in three independent experiments, and the standard deviation is indicated for each time point. **E**. Dodson kinetic test. Normalized product release curves for [Aurora B/IN-box]^loop-P^ (green) and [Aurora B/IN-box]^IN-P^ (magenta) at 25, 50 and 100 nM concentration in the presence of the peptide substrate and ATP. Note that the pink traces overlap during the first ∼50 minutes of incubation, indicating that phosphorylation of the activation loop is an intramolecular, concentration independent, reaction.

In [Aurora B/IN-box]^lN-P^, the phosphorylated TSS motif was rapidly stabilized by adjacent positively charged residues (IN-box^Lys839^, IN-box^Lys841^, and IN-box^Arg847^), albeit in a different conformation than in [Aurora B/IN-box]^all-P^ (Fig. S15B and movie M3). The phosphorylated IN-box interacts early in the simulation with the positively charged patch in the N-lobe of Aurora B and never extends to the C-lobe of Aurora B, in contrast to [Aurora B/IN-box]^all-P^. Therefore, in [Aurora B/IN-box]^lN-P^, the phosphorylated IN-box remains stably associated only with the N-lobe for the duration of the MD simulation. The stabilization of the N-terminal part of IN-box in this intermediate, that we observe with HDX, may be due to the interaction of the C-terminal IN-box with the N-terminal lobe of Aurora B. This interaction results in a more rigid IN-box, albeit in a conformation that is unproductive for enzyme activity.

A very interesting conformational change was observed in [Aurora B/IN-box]^loop-P^, where only the activation loop of Aurora B is phosphorylated, the C-terminal region of IN-box is immediately attracted to the Aurora B^pThr248^ (Fig. 5 and movie M4), in a similar manner as in [Aurora B/IN-box]^all-P^. In our simulation, we observe that Aurora B^pThr248^ retains the same hydrogen bonds as in the starting crystal structure of Aurora B (PDB 4C2W), in addition to Aurora B^Arg140^ also being recruited from the Aurora B^αC^-helix (Fig. 5B). Interestingly, an electrostatic interaction between Aurora B^pThr248^ and the IN-box^Arg843^ occurred after only 0.25 μs (which is even faster than a similar interaction in the case of [Aurora B/IN-box]^all-P^) and persisted throughout the rest of the simulation time (Fig. 5C). Finally, due to this large conformational change, the IN-box forms several other electrostatic and hydrophobic interactions along the N and C lobes of the enzyme. The final structure of [Aurora B/IN-box]^loop-P^ at the end of the simulation resembles [Aurora B/IN-box]^all-P^ (Fig. 2 and Fig. 5). This was a puzzling result, because both kinetics and HDX indicated significantly lower activity and a higher degree of disorder in [Aurora B/IN-box]^loop-P^ than in [Aurora B/IN-box]^all-P^. This unexpected result prompted us to perform a more detailed kinetic analysis.

### Phosphorylation of the activation loop is the intramolecular rate-limiting step in the activation process

Both partially phosphorylated complexes have low enzymatic activity when only one of the two sites is phosphorylated. They can recover their full enzymatic activity when the other site is phosphorylated during the autophosphorylation process (fig. S10A). We wanted to know whether one of the partially phosphorylated complexes can autoactivate faster than the other.

We preincubated [Aurora B/IN-box]^no-P^ (wt), [Aurora B/IN-box]^IN-P^, and [Aurora B/IN-box]^loop-P^ with ATP-Mg and monitored their autoactivation by determining the initial velocities after different preincubation times. We found that all three complexes can achieve the same activity as [Aurora B/IN-box]^all-P^ (wt) (Fig. S9A), but the autoactivation proceeds with different kinetics (Fig. 5D). As expected, the [Aurora B/IN-box]^no-P^, devoid of any phosphorylation, auto-activates slowest. However, the [Aurora B/IN-box]^loop-P^ is autoactivated much faster than the [Aurora B/IN-box]^IN-P^.

Previously, Aurora B autoactivation by phosphorylation was found to occur in two steps: first a slow intramolecular step followed by a faster intermolecular step (Zaytsev et al., 2016) but the tools to unambiguously correlate the kinetic steps with the specific phosphorylation events during autoactivation were not available. Our autoactivation experiments with partially phosphorylated intermediates clearly show that phosphorylation of the Aurora B activation loop is the slow step, as [Aurora B/IN-box]^IN-P^ (where Aurora B^Thr248^ is not phosphorylated) is slowly autoactivated. On the other hand, phosphorylation of IN-box is the fast step because [Aurora B/IN-box]^loop-P^ (where IN-box is not phosphorylated) autoactivates quickly. To further confirm that phosphorylation of the Aurora B activation loop occurs intramolecularly (cis) and phosphorylation of IN-box occurs intermolecularly (trans), we used the kinetic test developed by Dodson et al. (Dodson et al., 2013). To this end, we used different concentrations of enzyme complexes with fixed substrate concentrations and continuously followed the product formation (fig. S16A). The traces of product formation were normalized to the respective enzyme concentrations (Fig. 5E). An overlap of the normalized traces in the lag phase of the curve, indicates a concentration-independent, intramolecular, reaction. This profile was observed for [Aurora B/IN-box]^IN-P^ (pink curves), confirming that phosphorylation of the activation loop occurs in “cis.” In the case of [Aurora B/IN-box]^loop-P^ (green curves), the normalized traces of product formation at different enzyme concentrations show that the rate of product formation is concentration dependent. At higher enzyme concentration, the changes in the slope are more pronounced, confirming that phosphorylation of the TSS motif occurs intermolecularly. Thus, our kinetic analysis with partially phosphorylated intermediates identifies autophosphorylation of the Aurora B activation loop as a cis-acting, rate-limiting step and IN-box TSS phosphorylation as a faster trans-acting event in the autoactivation of [Aurora B/IN-box]. This mechanism has been suggested by Zaytsev, Segura-Peña et al (Zaytsev et al., 2016), but here we prove it experimentally, for the first time.

### The IN-box plays a crucial role in both organizing the substrate peptide binding site and enabling catalysis of the [Aurora B/IN-box] complex

According to the MD simulation the arginine residues of IN-box, IN-box^Arg843^ and IN-box^Arg847^, interact with Aurora B^pThr248^ in the intermediate [Aurora B/IN-box]^loop-P^ and the fully phosphorylated enzyme complex, respectively. To investigate the role of these arginine residues in kinase activity, we generated point mutants IN-box^Arg843Ala^ and IN-box^Arg847Ala^ in the context of the [Aurora B/IN-box] complex. Mutation of IN-box^Arg847^, which interacts with Aurora B^pThr248^ in the final structure of [Aurora B/IN-box]^all-P^, to alanine results in an enzyme complex with four times lower catalytic efficiency (v_max_/K_M_) (Table 1). This is due to the fivefold increase in K_M_ for the substrate peptide compared with the wild-type complex, while the k_cat_ increases only slightly. Thus, kinetic analysis identifies this residue and its interaction with Aurora B^pThr248^ as an important organizer of the substrate-peptide binding site. On the other hand, mutation of the IN-box^Arg843^ involved in stabilizing Aurora B^pThr248^ in [Aurora B/IN-box]^loop-P^ and stabilizing the IN-box TS^P^S^P^ conformation in [Aurora B/IN-box]^all-P^ simulations results in 2.5-fold decrease in catalytic efficiency. However, here the K_M_ for the substrate peptide is only slightly increased, while k_cat_ experiences a twofold decrease (Table 1), suggesting that the IN-box^Arg843^ predominantly supports the catalytic process. The residue is far from the active site of the enzyme complex but could be involved in stabilizing the enzyme conformations and/or dynamics, favoring efficient catalysis. These results again suggest a dual role of the IN-box residues (which is partially uncoupled in the tested mutants) in both substrate recognition and catalysis.

While analyzing the importance of the IN-box residues for catalytic efficiency, we also investigated the enzymatic activity of the double mutant IN-box^Lys846Asn-Arg847Gln^, in which two adjacent positively charged residues are replaced by neutral asparagine and glutamine. Originally, this mutant was generated with the idea of eliminating the Aurora B-substrate consensus site (positively charged residues preceding the serine or threonine to be phosphorylated (Alexander et al., 2011; Kettenbach et al., 2011)) at the IN-box, thereby preventing autophosphorylation of the IN-box TSS motif. Accordingly, this mutant was expected to have similar enzymatic activity as [Aurora B/IN-box]^loop-P^, but the double mutant [Aurora B/IN-box^Lys846Asn-Arg847Gln^] was virtually inactive, with a 400-fold lower k_cat_ compared with the wild-type enzyme (Table 1). Subsequent mass spectrometric analyzes showed that Aurora B^Thr248^ was not phosphorylated, in addition to the expected lack of phosphorylation of the TSS-motif of the IN-box. This observation underscores that the presence of positive charges at the IN-box is essential for the catalytic process and autoactivation. In addition to its function as a recognition sequence for autophosphorylation in the IN-box TSS -motif, these residues play an active role in stabilizing the productive enzyme complex.

### Maximal activation by phosphorylation is associated with concerted movement between the [Aurora B/IN-box] subdomains and the activation loop

Next, we analyzed the changes in global movements of the [Aurora B/IN-box] in MD simulations as a function of the phosphorylation status of the enzyme complex. We performed principal component analysis (PCA) for the MD trajectories of Aurora B in each of the phosphorylation states. The first principal components (eigenvectors) are always associated with the largest conformational changes, and this type of analysis has proven very useful for understanding global movements that govern catalysis in other protein kinases (Masterson et al., 2011). The dominant eigenvectors (fig. S17) for all enzyme forms describe the opening and closing of the active site and the twisting between kinase lobes, as well as the movements in the activation loop (Fig. 7A). However, the type of movement defined by the first principal component (PC1, the one corresponding to the largest relative movement) depends on the phosphorylation status of the enzyme complex. For [Aurora B/IN-box]^no-P^, PC1 is an open-close movement, for [Aurora B/IN-box]^IN-P^, it is a twisting, for [Aurora B/IN-box]^loop-P^, it is an activation loop movement, whereas for [Aurora B/IN-box]^all-P^, PC1 involves synchronized opening and closing of lobes and loop movement (Movie M5). Importantly, the cosine content evaluation (Hess, 2000) for PC1 in [Aurora B/IN-box]^all-P^ was only 0.01. In contrast, the first eigenvectors in all other forms had a cosine content between 0.5 and 0.9 (Fig. 7A, table). The small values of the cosine content indicate that the deformation in [Aurora B/IN-box]^all-P^ is a truly organized vibrational deformation that coordinates the open-closed/twist motion with the oscillation of the activation loop, whereas the first mode in all other phosphorylation states of the enzyme complex (with high cosine content) is mostly associated with random deformations.

We then analyzed the relationship between the open-closed movement and the twist movement (Fig. 7B). The plot shows that the unphosphorylated form of the enzyme complex, [Aurora B/IN-box]^no-P^ (blue), explores a wider range of conformations, consistent with a highly disorganized, easily deformable structure. Along the “open-closed” axis, this enzyme complex forms two groups of ensembles, clearly indicating the existence of two stabilized populations (one in the open and the other in the closed state; blue circles in the Fig. 6A), while intermediate conformations are sparse. Phosphorylation of the activation loop, [Aurora B/IN-box]^loop-P^ (green), significantly reduces the conformational space examined along both vectors. This reflects a better organized enzyme with closer distance between the lobes, consistent with progressive restriction of movement within the structure. Finally, full phosphorylation, [Aurora B/IN-box]^all-P^ (red), further increases the degree of order and rigidity within the enzymatic complex, forcing the enzyme into three well-defined regions in the graph that correspond to the occurrence of natural oscillations associated with opening and closing movements and twist. Negative twisting is associated with two distinct conformational basins, corresponding to the closed or open conformation of the enzyme. The separation of the two basins indicates that an open-closed transition is unlikely in this twist conformation. On the contrary, when the twist between the lobes is moderately positive, the enzyme can easily transition between the open and closed states. It can be concluded that in [Aurora B/IN-box]^all-P^, the transitions between open and closed conformation are regulated and coordinated by the twist, indicating complete synchronization of the global movements of the kinase subdomains when fully phosphorylated.

**Figure 6.**
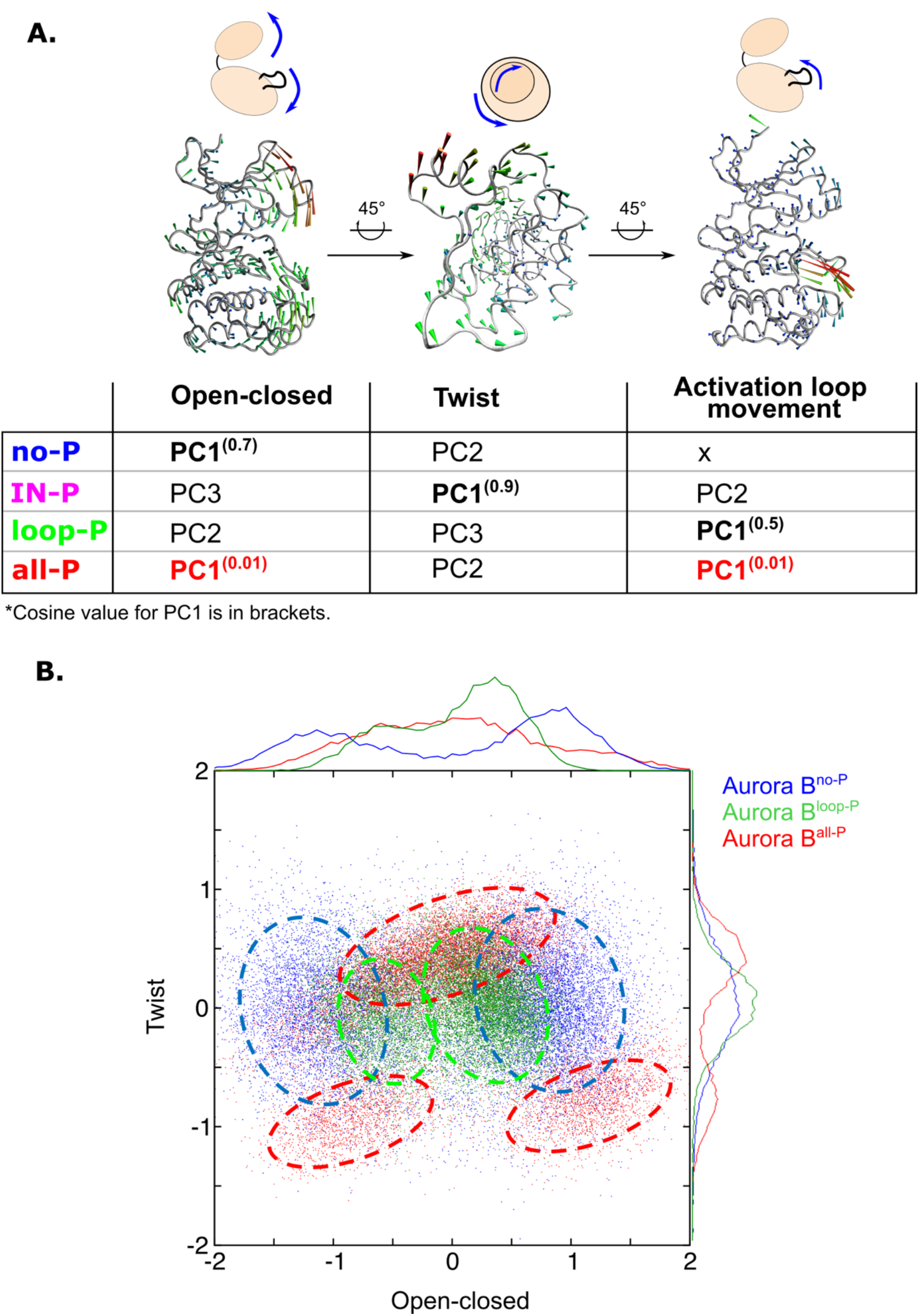
Principal component analysis (PCA) shows that only fully phosphorylated [Aurora B/IN-box] achieves coordinated movements between opening/closing, twisting and movements in the activation loop. **A**. Protein movements in the different phosphorylation forms of the [Aurora B/IN-box] complex in PC1 are illustrated by porcupine diagrams (opening/closing movement, twisting movement, movement in the activation loop). The needle tip indicates the direction of the vector and the magnitude, while the color (blue-green-red) corresponds to the increasing magnitude. The table summarizes the results of PCA for different phosphorylation forms of the complex. The cosine value close to 0, indicating an ordered movement, was obtained only for PC1 of [Aurora B/IN-box]^all-P^, implying a coordinated movement of opening and closing and a restructuring of the activation loop. **B**. Correlation between “opening/closing” and “twist” for [Aurora B/IN-box]^no-P^ (blue), [Aurora B/IN-box]^loop-P^ (green) and [Aurora B/IN-box]^all-P^ (red).

**Figure 7.**
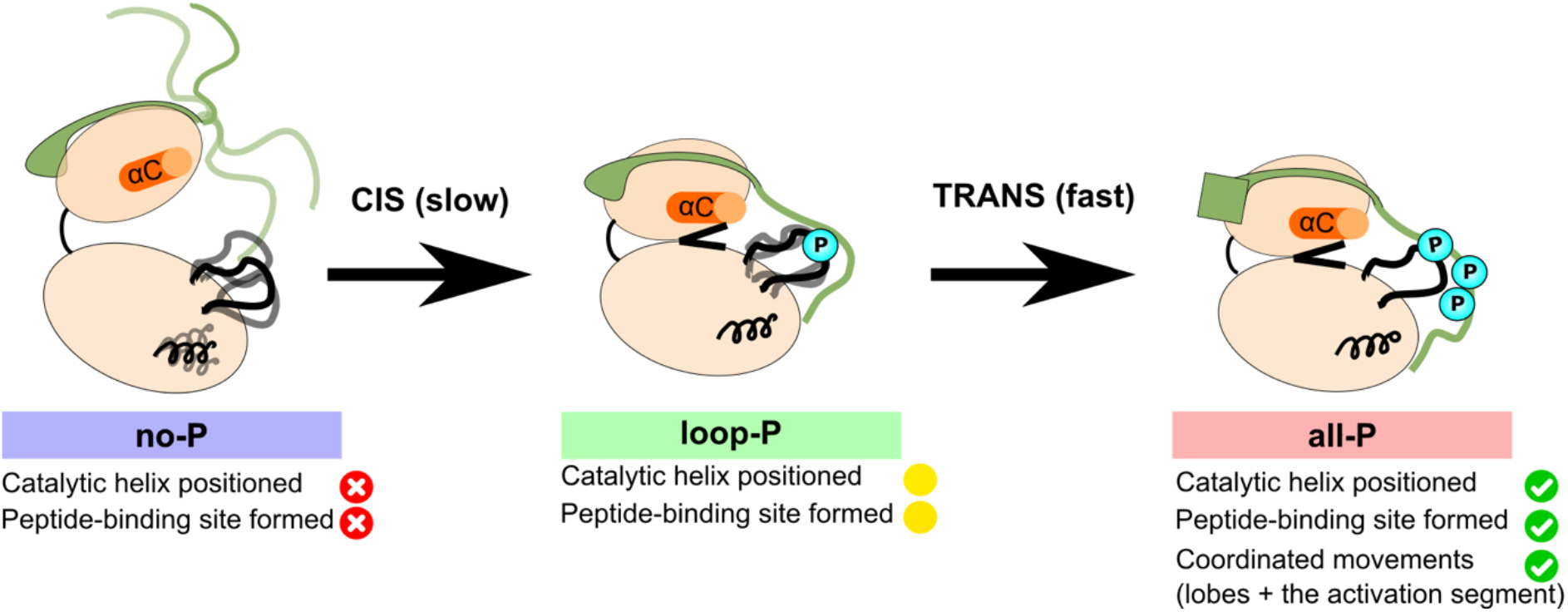
Model summarizing the autoactivation mechanism of [Aurora B/IN-box]. In the absence of phosphorylation, the [Aurora B/IN-box]^no-P^, the enzyme complex is very flexible. The activation segment and Aurora B^αG^ are disorganized, Aurora B^αC^ is not in catalytic position and the two lobes of the kinase are more separated. The IN-box is disorganized and its C-terminal region does not interact with Aurora B. This enzyme can autophosphorylate its own activation loop in a slow intramolecular process. Once the activation loop is phosphorylated, [Aurora B/IN-box]^loop-P^, Aurora B^αG^ is stabilized and the IN-box C-terminal region is recruited to interact with the C-lobe of Aurora B. This leads to the stabilization of Aurora B^αC^ and partial formation of the substrate binding site around the phosphorylated activation loop. This intermediate is further phosphorylated in the IN-box in a faster intermolecular process. In the fully phosphorylated form of the enzyme complex, [Aurora B/IN-box]^all-^ P, the IN-box stably interacts with the phosphorylated loop; Aurora B^αC^ is in the productive position and the peptide binding site is fully formed. The enzyme complex now has all functional elements in place and synchronization of key global motions of the kinase lobes ensures efficient catalysis.

In summary, only when the enzyme complex is fully phosphorylated are the opening and closing of the lobes, the rotation between the lobes, and the opening and closing of the activation loop fully coordinated, resulting in efficient catalysis. The PCA result is consistent with the kinetic and HDX analysis and highlights that full activity requires a well-structured enzyme complex with a high degree of coordination of movement, which is synergistically achieved by phosphorylation in the Aurora B activation loop and in the IN box.

## Discussion

Using hydrogen-deuterium exchange and molecular dynamics, we found that phosphorylation has a structuring effect on the [Aurora B/IN-box] complex, with the enzyme exploring fewer conformations and therefore becoming more rigid. Structuring after phosphorylation appears to be a general activation mechanism for the protein kinase family, and similar effects have been reported for protein kinase A and others (Hsu et al., 2008; Steichen et al., 2010).

A unique feature of Aurora B is its obligate association with the binding partner INCENP, which is also subject to phosphorylation. The recent crystal structure of [Aurora C/IN-box] showed that the phosphorylated IN-box interacts with the phosphorylated activation loop and stabilizes an active conformation of the kinase, implying a synergy between two phosphorylation sites (Abdul Azeez et al., 2019). We complement and extend these findings by using an experimental approach that is independent of the ability to crystallize the protein complex. Our approach addresses the lack of structural information for unphosphorylated complex and lack of information about the structural dynamics and its potential role in [Aurora B/IN-box] autoactivation. Using solution-based and computational approach we have methodically examined the unphosphorylated, phosphorylated, and partially phosphorylated forms of the enzyme complex under the same conditions. Our experiments reveal that the dynamics of the two partners (Aurora B and the IN-box) are linked and that phosphorylation of the activation loop in conjunction with phosphorylation in the IN-box is required to coordinate the dynamics between the N-terminal and C-terminal lobes of the kinase.

The HDX experiment clearly shows that the IN-box is a highly dynamic polypeptide chain, with most of its regions reaching maximal deuteration after only three minutes of D_2_O incubation (Data set D1), even in the phosphorylated state. Interestingly, the N-terminal part of the IN-box, the only part containing defined secondary structure elements, shows EX1-like HDX kinetics, suggesting that the IN-box^αA^ and IN-box^αB^ helices exist in an equilibrium between folded and unfolded states. Despite the unfolding of the helices, IN-box remains associated with Aurora B via many hydrophobic interactions (Fig. 2 and fig. S6), which is also confirmed by our mass photometry experiments of [Aurora B/IN-box]^no-P^. The EX1 kinetics in the N-terminal part of the IN-box is independent of the phosphorylation status but is more pronounced and faster in the unphosphorylated state. The former raises the possibility that in the unphosphorylated state, the unfolding event in the IN-box transfers disorder to the adjacent N-lobe of Aurora B and in this way negatively affects its kinase activity. Thus, in the unphosphorylated form of the enzyme complex, the IN-box acts as a negative regulator.

Oppositely, phosphorylation of the Aurora B activation loop results in partial structuring within Aurora B and a major conformational change of the IN-box as well as in adoption of an enzyme conformation close to that required for efficient catalysis. However, the lack of phosphate groups on the IN-box continues to interfere with the optimal arrangement of the enzyme complex. Finally, when both Aurora B and the IN-box are phosphorylated, the enzyme complex is structured and coordinated in its global motions. Thus, in the fully phosphorylated form of the enzyme complex, the phosphorylated IN-box acts as a positive regulator that stabilizes the productive enzyme conformation. The results from HDX and MD simulation suggest that the dynamics of Aurora B and the IN-box are coupled in their interacting regions.

It is noteworthy that the HDX method used here can detect differences in H/D exchange of the protein backbone on a time scale of seconds to hours, whereas changes on a faster time scale are not detectable with our setup. This may be the reason why our experiments did not detect phosphorylation-induced changes in very flexible parts of the protein, such as the activation loop of Aurora B and the C-terminus of the IN-box. For peptides covering these regions, we see that complete exchange occurs at very early time points (Data set D1: peptides covering 231-244 - activation loop and peptides covering 821-864 - C-terminus of IN-box). To fully capture the dynamics of the enzyme complex in these regions, one would need to use approaches that can capture the dynamics at lower time scales, such as stopped-flow time-resolved HDX (Al-Naqshabandi and Weis, 2017) or NMR (nuclear magnetic resonance).

Using a chemical ligation approach, we generated partially phosphorylated intermediates that allowed us, for the first time, to describe the role of each of the major phosphorylation sites in the enzymatic activity and dynamics of [Aurora B/IN-box] (Fig. 7). Our kinetic, HDX, and MD experiments with partially phosphorylated intermediates confirmed that synergy between two phosphorylation sites is necessary not only for the adoption of the active conformation of the enzyme complex (already indicated in the crystal structure of fully phosphorylated [Aurora C/IN-box] (Abdul Azeez et al., 2019) but also for the achievement of full enzymatic activity (Fig. 3C), full structural stability (Fig. 4), and coordination between movements within the enzyme complex (Fig. 6).

It has been previously proposed that autoactivation of [Aurora B/IN-box] occurs via a two-step mechanism involving a slow cis-step and a fast trans-step (Zaytsev et al., 2016). Following our kinetic and MD analysis with partially phosphorylated enzyme complexes, we can now confirm this model and conclude that phosphorylation of the Aurora B activation loop is a slow intramolecular process followed by the fast trans phosphorylation of the IN-box TSS motif.

Phosphorylation gradients of Aurora B substrates have been identified in the cell (Fuller et al., 2008; Wang et al., 2011) and these long-range gradients play a regulatory role in mitosis (Afonso et al., 2014; Ferreira et al., 2013; Uehara et al., 2013). In addition, a strong evidence for the existence of steep gradient of Aurora B activity at the metaphase chromosomes has been reported and it has been proposed that the kinetics of Aurora B activity plays an important role in the formation of this gradient (Liu et al., 2009; Zaytsev et al., 2016). Here, we found that the partially phosphorylated [Aurora B/IN-box] intermediates have very low kinase activity (5-10%), which, together with the two-step Aurora B autoactivation, is consistent with the ability to form a steep phosphorylation gradient. For example, a small increase in phosphatase concentration in a local cellular region like the kinetochore, could produce a large decrease of kinase activity in the kinetochore region, while in a nearby region the Aurora B kinase activity could remain high. In addition, the fast autoactivation kinetics of Aurora B observed with the [Aurora B/IN-box]^loop-P^ intermediate is consistent with the *in vivo* observation of waves of Aurora B kinase activity. These waves start in a site of Aurora B clustering and rapidly propagate to distant regions with lower concentrations of Aurora B (Zaytsev et al., 2016).

At the structural level, upon phosphorylation of the IN-box TSS motif, an optimal interaction between the IN-box C-terminal region and the activation loop take place. Only in this optimal conformation, when two intrinsically disordered parts, the Aurora B activation loop and the C-terminus of the IN-box, interact, do allosteric changes confer the highest degree of order to the enzyme complex and movements between the kinase lobes become synchronized, resulting in full enzymatic activity.

It has been previously reported that domain movements and correlated motions within kinase lobes are tightly coupled to enzymatic turnover (Kumar et al., 2018; Kuzmanic et al., 2017; Masterson et al., 2010; Xiao et al., 2014). The strongest evidence for the coupling of global dynamics of protein kinase lobes with catalysis comes from NMR experiments on Protein kinase A (PKA), where the opening and closing of the active site correlates very well with the catalytic turnover of the enzyme (Masterson et al., 2011). Since only the fully active [Aurora B/IN-box]^all-P^ exhibits synchronized global motions, we propose that global domain movement is directly linked to the catalytic cycle of this enzyme complex and likely controls substrate binding and product release.

The very similar Aurora A (72% identity), which associates with a different binding partner, TPX2, can be activated to an appreciable extent independently, either only by association with TPX2 or only by phosphorylation in the activation loop (Dodson and Bayliss, 2012). Two pathways of Aurora A activation are used separately at the physiological level. The pool of Aurora A involved in chromatin-driven microtubule nucleation is regulated exclusively by association with TPX2 (Kufer et al., 2002) and phosphorylation in the activation loop is not observed (Levinson, 2018). On the other hand, Aurora A activation at the centrosome depends only on phosphorylation of the activation loop and does not involve TPX2 binding (Joukov et al., 2010). While Aurora A has evolved to allow two independent activation pathways, activation of Aurora B, which is always in complex with IN-box, follows a single synergistic activation pathway that requires the coincidence of two events: Phosphorylation of threonine in the activation loop and phosphorylation of the binding partner (TSS motif in the IN-box).

The present study demonstrates the important role that dynamics plays in regulating the enzymatic activity of [Aurora B/IN-box]. Our study explains and highlights the importance of IN-box (specific and obligatory Aurora B partner) as a necessary structural and strong regulatory element of the enzyme complex. The new findings suggest that the interface between Aurora B and the IN-box represents a target area for the development of a new type of Aurora B-specific drugs that would disrupt the allosteric changes required for activation. Our results contribute to a better understanding of the biophysical basis for the switch-like transition in Aurora B catalytic activity upon autophosphorylation (Zaytsev et al., 2016), a feature that underlies the intracellular spatial pattern of Aurora B substrate phosphorylation essential for proper chromosome segregation and cell division.

## Materials and methods

### Preparation of [Aurora B/IN-box] complex in different phosphorylation states

The wild type (wt) enzymatic complex containing Aurora B (60-361 aa) and IN-box (790-856 aa) from *Xenopus laevis* was expressed in *E. coli*, using a bicistronic construct that was cloned in the pRSF duet vector (Novagen). The GB1 domain from the streptococcal protein G was fused to the N-terminus of Aurora B in order to increase protein expression and solubility. BL21(DE3)pLysS cells were transformed with the mentioned plasmid and protein expression was induced with 0.2 mM of IPTG at OD of 0.6 and incubated for 18 hours at 18 °C with shaking at 180 rpm.

Protein purification was conducted in two steps. First, IMAC with Nickel column (GE healthcare) was used to trap the tagged protein complex. After removal of the 6XHisGB1 by TEV cleavage, the enzyme complex was further purified with size exclusion chromatography. The protein was stored at -80 °C in a buffer containing 10 mM Hepes pH 7.5, 150 mM NaCl, 2.5 mM DTT and 50 % glycerol until further use. To obtain the dephosphorylated form, the enzyme complex was incubated with 0.2 μM of GST-λ phage phosphatase (made in-house) during one hour at 30 °C, followed by incubation with glutathione beads to remove the phosphatase.

To generate the partially phosphorylated forms of the enzyme, the 6XHis-GB1-Aurora B (59-361 aa) and IN-box (790-844 aa; IN-box^ΔC^) coding regions were introduced in the pTXB1 plasmid (NEB) by in-fusion cloning using the Nde1 and Sap1 cloning site. The resulting construct has two translational cassettes. The upstream cassette contains 6xHis-GB1-(TEV site)-Aurora B and the downstream cassette contains the IN-box^ΔC^ fused to the N-terminus of the Mxe intein-chitin binding domain protein (Telenti et al., 1997). The enzyme complex, [Aurora B/IN-box]^IN-ΔC^, was expressed using the same parameters as the WT. Furthermore, IMAC was used as the first step of purification. Next, for obtaining the [Aurora B/IN-box]^loop-P^, the enzyme complex was incubated with the fully active [Aurora B/IN-box] (10 nM final) in the presence of 5mM ATP-Mg for 1 hr. This ensured full phosphorylation of Aurora B^Thr248^. Analogously, for obtaining [Aurora B/IN-box]^IN-P^, the enzyme complex was incubated with GST-λ-phosphatase (0.2 μM final) for 1 hr. This ensured full dephosphorylation of Aurora B^Thr248^. In parallel with either phosphorylation or dephosphorylation of the enzyme complex, the reaction was incubated with TEV protease for 3 hours to remove the 6xHis-GB1 tag. The enzyme complex was then further purified by incubation with chitin beads (NEB), to remove traces of TEV and fully active kinase or phosphatase. Elution from the chitin beads was done by overnight incubation with 50 mM of 2-mercaptoethanesulfonic acid (MESNA) (Sigma Aldrich) that induces intein-mediated cleavage. The product is a C-terminal thioester on the IN-box^ΔC^ which can be ligated to a synthetic peptide with an N-terminal cysteine. At this step, in order to obtain [Aurora B/IN-box]^loop-P^, we fully incubated phosphorylated [Aurora B/IN-box]^IN-ΔC^ with the peptide sequence CKRTSSAVWHSPPL-6xHis. Analogously, in order to obtain [Aurora B/IN-box]^IN-P^ we incubated dephosphorylated [Aurora B/IN-box]^IN-ΔC^ with the peptide sequence CKRTS(PO_4_)S(PO_4_)AVWHSPPL-6xHis. The ligation reaction proceeded overnight in a ligation buffer (50 mM HEPES pH 8.0, 300 mM NaCl, 10 mM MESNA, 5 mM Ascorbic acid, 10 mM TCEP) with 50 μM [Aurora B/IN-box]^ΔC^ and 600 μM peptide. To remove the excess of unligated peptide, the protein sample was injected un Superdex S200 16/60 column. To separate the ligated enzymatic complex from the un-ligated [Aurora B/IN-box]^IN-ΔC^, the sample was run over a Ni column and the pure ligated protein was eluted with imidazole. The protein was stored same as WT until further use.

### Enzymatic assay

The enzymatic activity of Aurora B was measured by monitoring the product (ADP) formation rate using a pyruvate kinase/lactate dehydrogenase-coupled spectrophotometric assay (Roskoski, 1983; Wu and Wang, 2003). The peptide substrate used was ALRRFSLHGA, which was acetylated at the N terminus and amidated at the C terminus. Kinetic measurements were performed in kinase assay buffer (25 mM Tris-HCl pH 7.4, 100 mM KCl, 2 mM MgCl_2_, 1 mM EGTA, 2 mM DTT, 1 mM ATP, 160 μM NADH, 0.5 mM phosphoenol pyruvate containing 20 U/ml lactate dehydrogenase and 20 U/ml pyruvate kinase) with substrate peptide at the concentrations indicated in the experiment. The reaction was initiated by addition of the [Aurora B/IN-box] complex, and the progress of the reaction was continuously monitored by tracking the decrease in absorbance at 340 nm using a Cary UV 60 spectrophotometer (Agilent). The initial rate was determined by calculating the slope of product release in the linear portion of the reaction, which occurs in the first few minutes of the reaction when less than 10% of the substrate has been converted to products.

### Aurora B autoactivation assay

The autoactivation of [Aurora B/IN-box]^no-P^, [Aurora B/IN-box]^loop-P^ or [Aurora B/IN-box]^IN-P^ was assessed by preincubating 12 μM of the enzymatic complex in a buffer containing 20 mM Tris pH 7.4, 100 mM KCl and 5 mM ATP-Mg. The autophosphorylation reaction was left to proceed and aliquots were taken at different times of preincubation with ATP to determine the initial rates. Each aliquot was diluted to a low nanomolar concentration (10 to 50 nM in the kinase assay buffer supplemented with 0.6 mM substrate peptide) and initial rates were determined as described above. Since low nanomolar concentrations of Aurora B were used in the assay and the rate is determined in the first minutes of the reaction (2-4 min; where the kinetics of product release are linear), the amount of autophosphorylation of the enzyme is negligible.

### Kinetic test to determine cis or trans activation steps

In difference to the autoactivation assay, where enzyme complex was pre-incubated with ATP-Mg, here the enzyme complexes at different concentrations (25 nM, 50 nM, and 100 nM) were incubated with 1 mM ATP-Mg and 0.6 mM substrate peptide at the same time, and the formation of ADP was continuously monitored for a prolonged period (450 min). It should be noted that the traces of product formation are not linear due to the autoactivation that is happening during the experiment. The traces of ADP formation are normalized to the concentration of the enzyme complex to evaluate the concentration effect on the autoactivation process, as reported by Dodson et al. (42). Overlapping curves in the lag phase of the product formation indicate concentration-independent activation (cis) and curves with different slopes indicate concentration-dependent activation (trans).

### Hydrogen deuterium exchange assay

The on-exchange reaction was done by diluting [Aurora B/IN-box] enzymatic complex 1:20 in a the on-exchange buffer (20 mM HEPES pD 7.0, 300 mM NaCl, 2mM DTT, 1 mM ADP, and 2 mM MgCl_2_, in D_2_O) so that the final D_2_O concentration was 95%. The on-exchange reaction was quenched at different time points (3, 10, 30, 90 and 300 minutes) by mixing 50 μl of the reaction with equal volume of ice-cold quench buffer (250 mM potassium phosphate pD 2.3) and flash freezing in liquid nitrogen. The quenched samples were stored at -80°C before proteolysis and liquid chromatography–mass spectrometry steps. The HDX experiment was performed with three independently prepared samples (biological replicates) for the wild type [Aurora B/IN-box]^all-P^ and [Aurora B/IN-box]^no-P^ and one one biological replicate for the intermediates phosphorylated forms. All the biological replicates show identical results. The reported data is for one biological replicate where each time point was run in duplicate or triplicate.

### Liquid Chromatography and Mass Spectrometry

The quenched samples were thawed on ice and injected on a nanoACQUITY UPLC system with HDX technology (Waters). The temperature of the chamber containing the sample loop as well as the UPLC and trap columns was set at 0.5°C. The temperature of the pepsin column compartment was set at 10 oC. The quenched samples (10 picomol) were injected into a 50 μl sample loop and run in a trapping mode, where the protein was passed through a pepsin column (Waters Enzymate 2.1 × 30 mm, 5 μm) and the proteolyzed sample was immediately directed to a trap column (Waters Acquity Vanguard BEH C18,1.7 μm, 2.1X 5 mm) to desalt peptides fragments. The flow rate was set to 70 μl/min during the first minute, followed by 100 μl/min for another 2 minutes with buffer A (0.2% formic Acid, 0.01 % Trifluoroacetic Acid pH 2.5). After desalting, the peptides were separated by C18 analytical column (Waters Acquity BEH C18, 1.7 μm, 1.0 × 100 mm) with a linear 5–50% acetonitrile gradient using buffer B (99.9% acetonitrile, 0.1% formic acid and 0.01 % Trifluoroacetic Acid pH 2.5). The elution gradient was run at 40 μl/ min for 17 min. The output of the analytical column was directed to a mass spectrometer (Q-TOF SYNAPT G2-Si, Waters) for peptide identification and determination of the deuterium uptake. The mass spectrometer was operated in the positive ion electrospray mode, with the ion mobility function to minimize spectral overlap using the MS^E^ acquisition mode (Waters Corporation). Lock mass correction with the Leu-ENK peptide was used to ensure mass accuracy determination.

To prepare fully deuterated sample, the unlabeled enzyme complex was injected to the HPLC system for digestion on pepsin column under the same condition as other samples. However, the peptides eluted from the analytical column were collected in one fraction instead of being injected to MS. The peptides were then lyophilized and resuspended in the on-exchange buffer and the reaction was injected into the HPLC system connected to MS in the same way it was for the other samples but bypassing the pepsin column.

### HDX-MS data evaluation

A library of non-deuterated peptides was created using the ProteinLynx Global server 3.0 (PLGS) (Waters) using the following requirements: 1.) A mass error for the peptide has to be below 10 ppm for the precursor ion, 2.) The peptide has to have at least two fragmentation products and 3.) The peptide has to be identified in at least 75% of the non-deuterated runs, where at least three replicates were used. The level of deuteration in the peptides was determined with DynamX 3.0 (Waters). A manual inspection of all the assignment was conducted to confirm the data or discard the noisy or overlapping spectra.

The percentage of deuteration for wild-type [Aurora-B/IN-box]^all-P^ and wild-type [Aurora-B/IN-box]^no-P^ was calculated by normalization with respect to the theoretical maximum uptake (MaxUptake), which was set at 100%. The maximum uptake is determined as follows: MaxUptake= N-P-2, where N is the number of amino acids in the peptide and P is the number of prolines. The percentage of deuteration was determined according to formula: 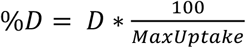.

The percentage of deuteration for [Aurora-B/IN-box]^loop-P^, [Aurora-B/IN-box]^IN -P^, [Aurora-B/IN -box]^all-^ P*, and [Aurora-B/IN-box]^no-P*^ was calculated by normalization with respect to the deuterium uptake in the fully deuterated peptides, which was set at 100%. The percentage of deuteration was determined by the following formula: 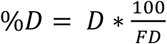where D is the amount of deuterium for a peptide at a given time point and FD is the amount of deuterium determined in the fully deuterated peptide. The [Aurora-B/IN-box]^all-P*^ and [Aurora-B/IN-box]^no-P*^ constructs contain the mutation IN-box^Phe845Cys^ to correspond to the partially phosphorylated intermediates where this mutation is present due to the protein ligation process.

All peptides with a deuteration difference greater than 5% were considered different.

### Mass photometry assay

The protein sample [Aurora-B/IN-box]^all-P^ and [Aurora-B/IN-box]^no-P^ were kept at 100 nM concentration in buffer containing 300 mM NaCl, HEPES 25 mM pH 7.5 and 2 mM DTT. Immediately before the measurement the sample was diluted to 10 nM. The data was acquired with Refeyn Two MP and analyzed with the Refeyn Discover MP software.

### Molecular dynamics simulations

We modelled the [Aurora B/IN-box] complex in four distinct phosphorylation states (no-P, loop-P, IN-P, all-P). Every model was independently built using the x-ray structure of the [Aurora B/IN-box]^loop-P^ complex from *X. laevis* (PDB ID:4C2W (Sessa and Villa, 2014). The non-hydrolysable adenylyl-imidodiphosphate ATP mimic present in the original crystal structure was replaced with the ATP-Mg^2+^ moiety taken from Aurora A structure (PDB ID:5DN3;(Janeček et al., 2016). The ^848^TSSAVWHSP^856^ C-terminal region of IN-box, not visible in the original crystal structure, was modelled as a random coil. The initial structures of the three phosphorylation states for which an experimental structure is not available were modelled by introducing of the pertinent phosphorylated groups where needed or removing the one in Aurora B^Thr248^. The starting structures were completed by the addition of the missing hydrogens, choosing standard protonation at neutral pH for all titratable groups. Each of the four systems were solvated by around 36’000 water molecules and contained in a periodic box with an edge of 105 Å. The systems were completed by the addition of 113 Cl^-^ and the corresponding Na^+^ required to achieve charge neutrality, resulting in a salt concentration close to the physiological value of 150 mM.

The CHARMM36 force-field (Huang et al., 2017) was used to parameterize the protein segments, ATP, and the ions, while water was represented with the triangular TIP3P model (Jorgensen et al., 1983) All bonds involving hydrogen atoms were constrained to their equilibrium distances using the LINCS algorithm (Hess et al., 1997), the Lennard-Jones potentials were computed using a cut-off distance of 10 Å, while the electrostatic interactions were evaluated using the Particle-Mesh Ewald method (Essmann et al., 1995).

The four systems were first relaxed to a low energy state by the steepest-descent algorithm until reaching a tolerance value of 1000 kJ mol^-1^ nm^-1^ on all forces. Then, the system was by molecular dynamics in the NPT ensemble. The equations of motion were integrated using the leap-frog algorithm (Hockney R. (last) and Eastwood J. W., n.d.) with a timestep of 2 fs. Pressure coupling was obtained using a Berendsen barostat, with coupling constant of 2 ps, and target pressure 1 Bar. Temperature equilibration was obtained by initially randomly setting the velocities of all atoms using a Maxwell-Boltzmann distribution at 10 K, and then by performing simulated annealing to 300 K within 2 ns of simulations. After equilibration, each system underwent a production run of 2.8 μs in the NPT ensemble, using the Nosé-Hoover chain thermostat (Martyna et al., 1992; Nosé, 1984; Parrinello and Rahman, 1981) with a coupling constant of 1 and 2 ps, respectively.

All simulations were performed and partly analyzed using the GROMACS package (Pronk et al., 2013; Van Der Spoel et al., 2005). Additional analysis and visualization were done using VMD (Humphrey et al., 1996).

## Acknowledgments

The authors acknowledge the support of the Research Council of Norway through the Centre for Molecular Medicine Norway (Project No. 187615; N.S., S.M.H.W. and D.S.P.), CoE Hylleraas Centre for Quantum Molecular Sciences (Project No. 262695; O.H. and M.C.), the Norwegian Supercomputing Program (NOTUR) (Project No. NN4654K; O.H. and M.C.) and individual Research Council of Norway grant (Project No. 325528) to N.S. The work in the Black lab was supported by the NIH: Grant No. R35-GM130302 to B.E.B. J.M.D.-M. was supported by NIH postdoctoral fellowship GM108360. We would also like to thank Prof. Magnus Kjærgaard (Århus University, Denmark) for performing the mass photometry measurements.

D.S.P., N.S. and B.E.B conceived the project. D.S.P carried out cloning, protein expression and purification, peptide design and ligation, kinetic characterization, HDX experiments and HDX data processing and analysis. O.H. performed MD experiments under supervision of Mi.C. O.H., Ma.C. and Mi.C. analyzed and interpreted MD data. J.M.D.-M. performed initial HDX experiments, designed and purified proteins, and performed initial phosphopeptide identification and quantitation under the direction of B.E.B. H.G. and S.M.H.W. helped with protein purification. D.S.P. and N.S. supervised the project and wrote the manuscript with input from all authors.

The authors declare that they have no conflict of interest.

All data needed to evaluate the conclusions in the paper are present in the paper and/or the Supplementary Materials.

