## Supplementary material for "The structural basis of the multi-step allosteric activation of Aurora B kinase"

**This PDF file includes:**

Figs. S1 to S17
Data set D1

**Other Supplementary Materials for this manuscript include the following:**

Movies M1 to M5

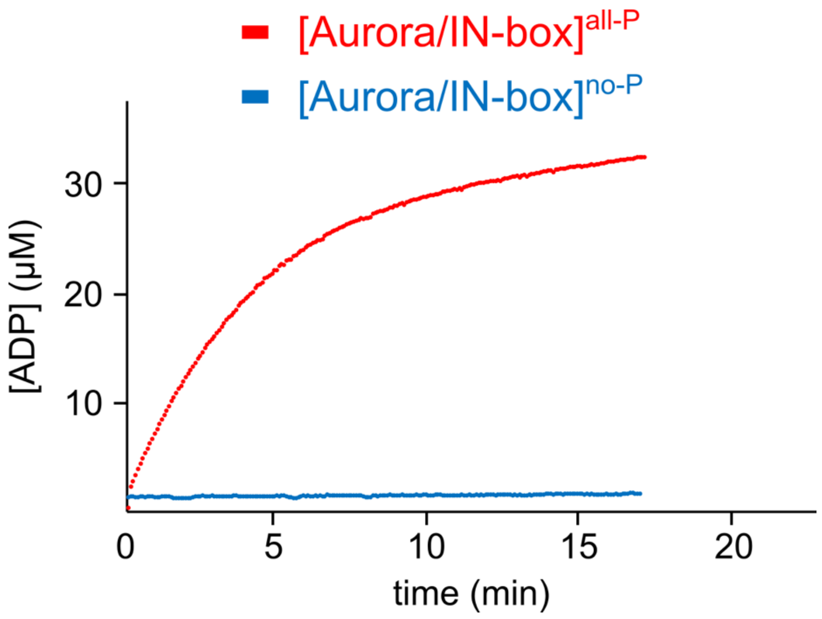


**Fig. S1.**

Time course of product formation. [Aurora B/IN-box]$^{no\text{-}P}$ (blue) and [Aurora B/IN-box]$^{all\text{-}P}$ (red). The [Aurora B/IN-box]$^{no\text{-}P}$ and [Aurora B/IN-box]$^{all\text{-}P}$ concentration in the assay was identical (20 nM).

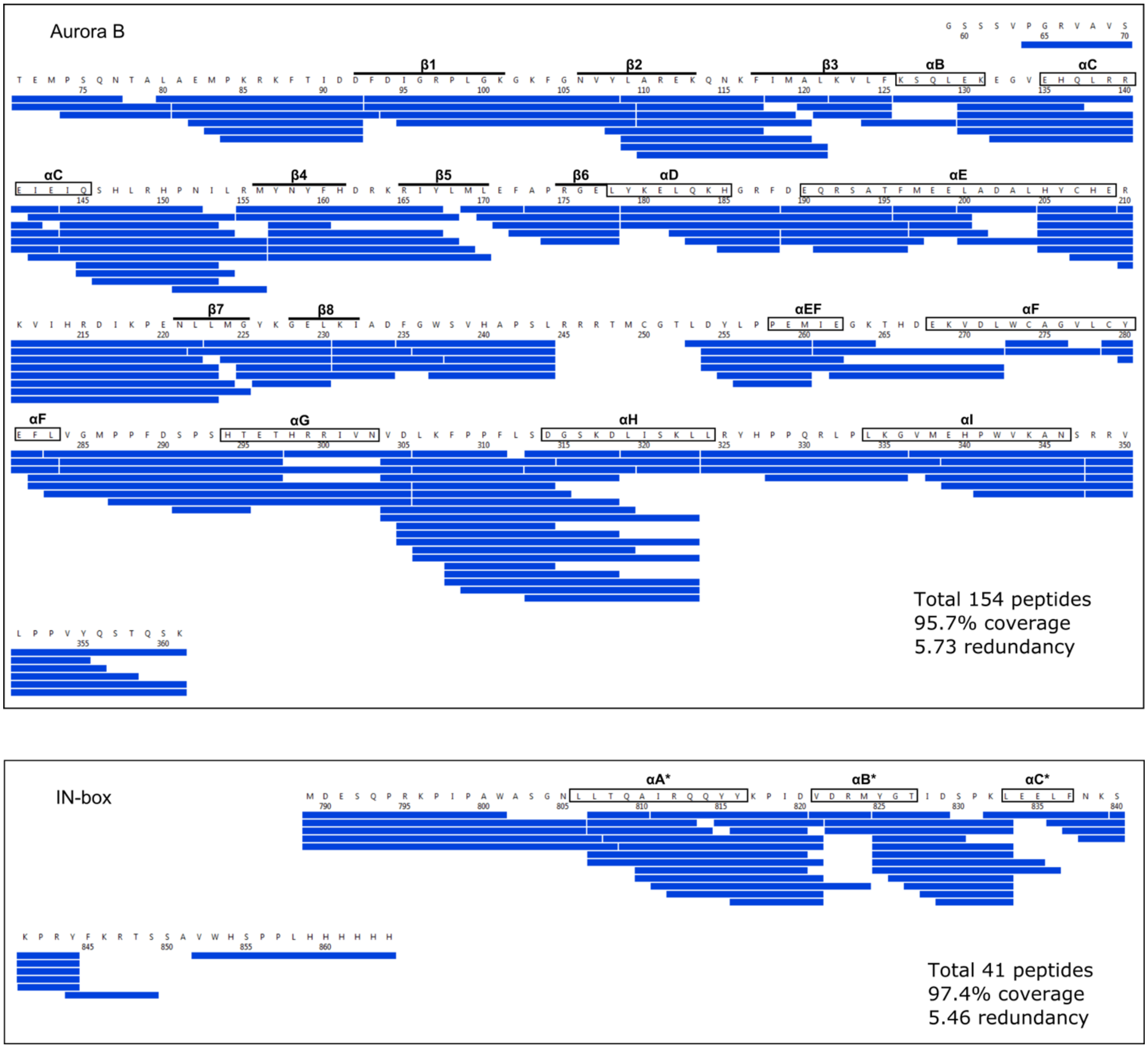


**Fig. S2.**

The HDX peptide coverage map for Aurora B and IN-box in HDX experiments. Each blue rectangle represents a unique peptide.

### IN-box

$^{807}$LTQAIRQQYYKPIDV$^{821}$

#### [Aurora/IN-box]$^{no-P}$

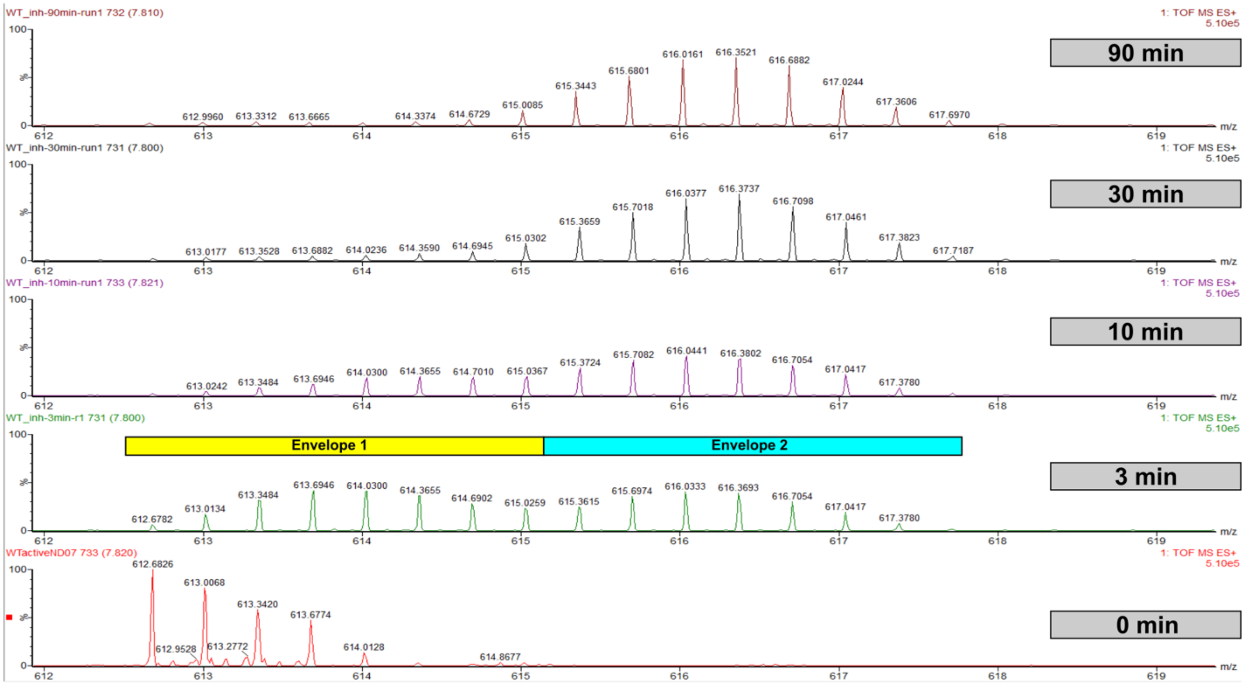


#### [Aurora/IN-box]$^{all-P}$

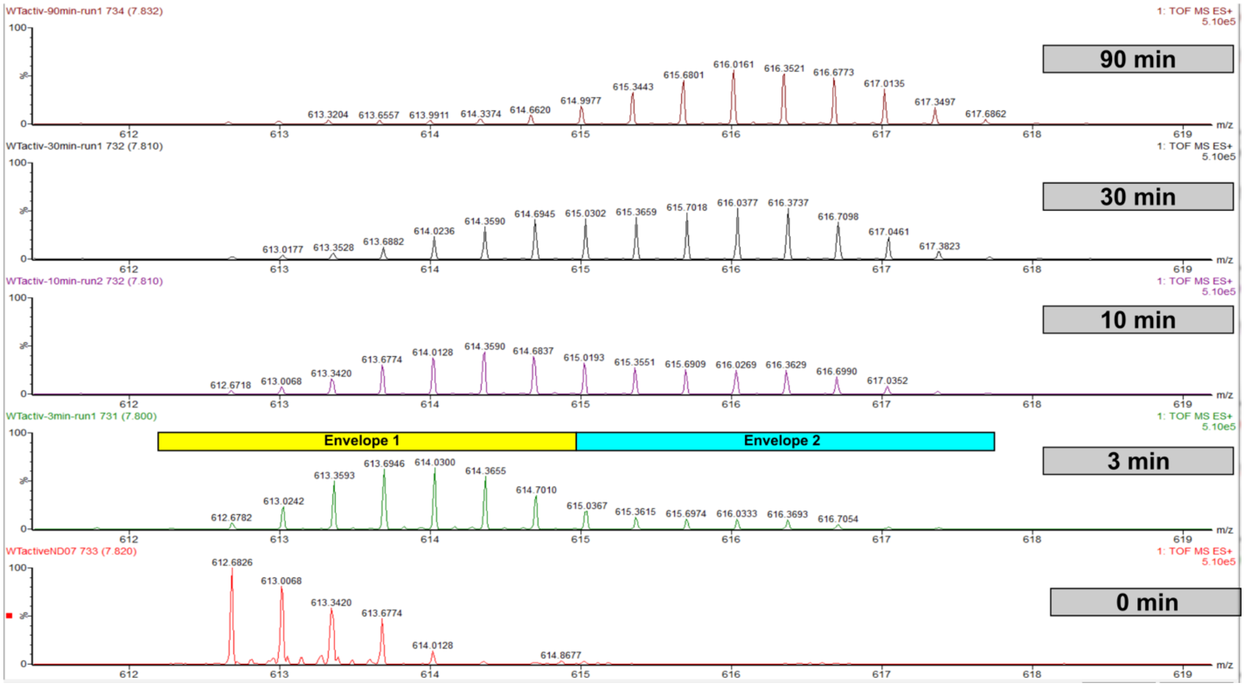

**Fig. S3.**

EX1 kinetics observed in IN-box$^{\alpha A}$, peptide $^{807}$LTQAIRQQYYKPIDV$^{821}$, in [Aurora B/IN-box]$^{no\text{-}P}$ and [Aurora B/IN-box]$^{all\text{-}P}$. The horizontal rectangles in the 3 minutes deuterated sample indicate the bimodal distribution of deuteration. The yellow rectangle is above the envelope with the lower m/z center of mass, representing molecules in which the IN-box$^{\alpha A}$ undergoes slow H/D exchange (HDX) and the cyan rectangle is above the envelope with the higher M/Z center of mass, where HDX occurs simultaneously throughout the helix due to an unfolding event. Note that at the 3-minute time point, the population with the higher m/z center is equal to the population with lower m/z center in [Aurora B/IN-box]$^{no\text{-}P}$, whereas in [Aurora B/IN-box]$^{all\text{-}P}$ the population with the lower m/z center of mass dominates, indicating a more stable IN-box$^{\alpha A}$ in [Aurora B/IN-box]$^{all\text{-}P}$.

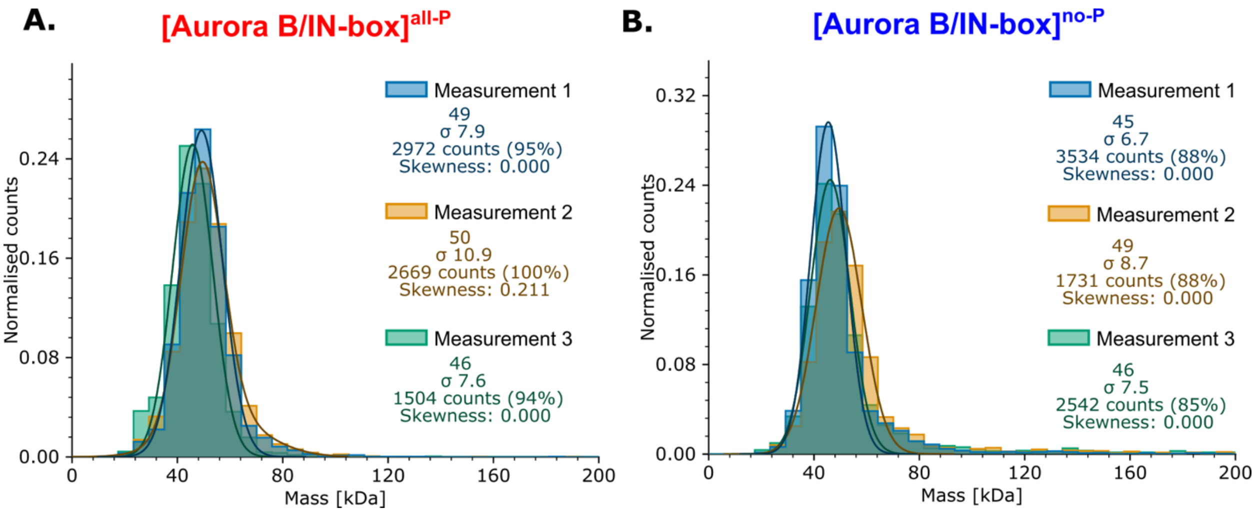


**Fig. S4.**

Mass photometry analysis of [Aurora B/IN-box]$^{all-P}$ (A) and [Aurora B/IN-box]$^{no-P}$ (B).

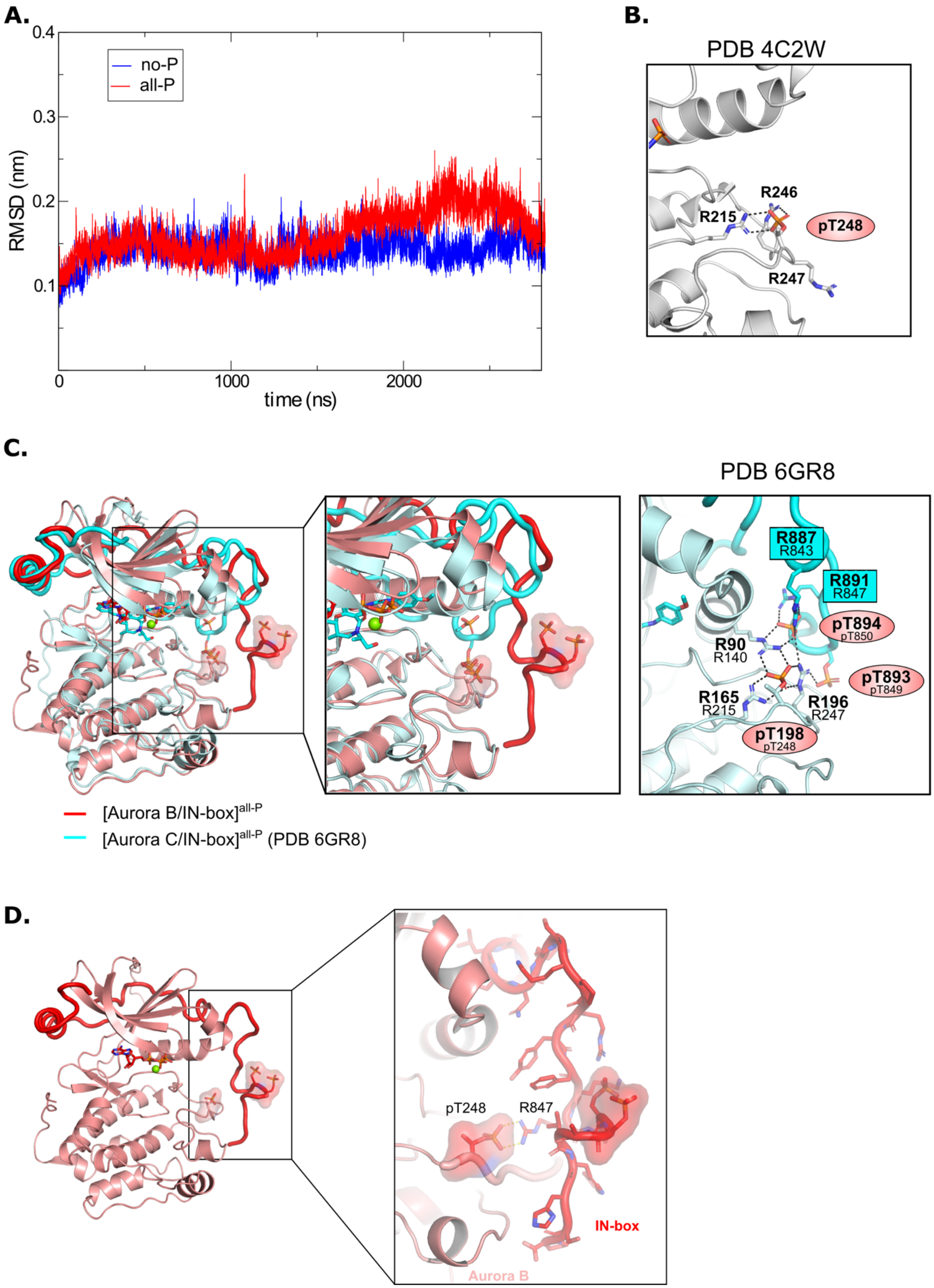
A.
0.4
0.3
0.2
0.1
RMSD (nm)
no-P
all-P
0
1000
2000
time (ns)
B.
PDB 4C2W
R246
R215
pT248
R247
C.
PDB 6GR8
R887
R843
R891
R847
pT894
pT850
R90
R140
pT893
pT849
R165
R215
R196
R247
pT198
pT248
[Aurora B/IN-box]all-P
[Aurora C/IN-box]all-P (PDB 6GR8)
D.
pT248
R847
IN-box
Aurora B

**Fig. S5.**

**A.** Root-mean-square-deviation (RMSD) for [Aurora B/IN-box]$^{no\text{-}P}$ (blue) and [Aurora B/IN-box]$^{all\text{-}P}$ (red) complexes during 2800 ns MD simulations. **B.** Zoomed view of the hydrogen bonding network around Aurora B$^{pThr248}$ as seen in the initial crystal structure (PDB 4C2W). **C.** (Left) Best overall overlay between MD model from our simulation, [Aurora B/IN-box]$^{all\text{-}P}$ (red), and the crystal structure of phosphorylated human [Aurora C/IN-box] (cyan). (Middle) Zoom views of the activation loop illustrating similarities/differences between the two structures. (Right) Hydrogen bonding network around Aurora B$^{pThr198}$ (analogous to Aurora B$^{pThr248}$ in *X. laevis* Aurora B) observed in the crystal structure of phosphorylated human [Aurora C/IN-box]. Numbers of residues corresponding to the X. laevis construct are given below the numbers in the human complex. **D.** Magnified view of the C-terminal part of the IN-box in the final MD conformation of [Aurora B/IN-box]$^{all\text{-}P}$ showing that despite the Aurora B$^{p248}$-IN-box$^{Arg847}$ electrostatic interaction, the main chain hydrogen atoms are available for H/D exchange with the solvent.

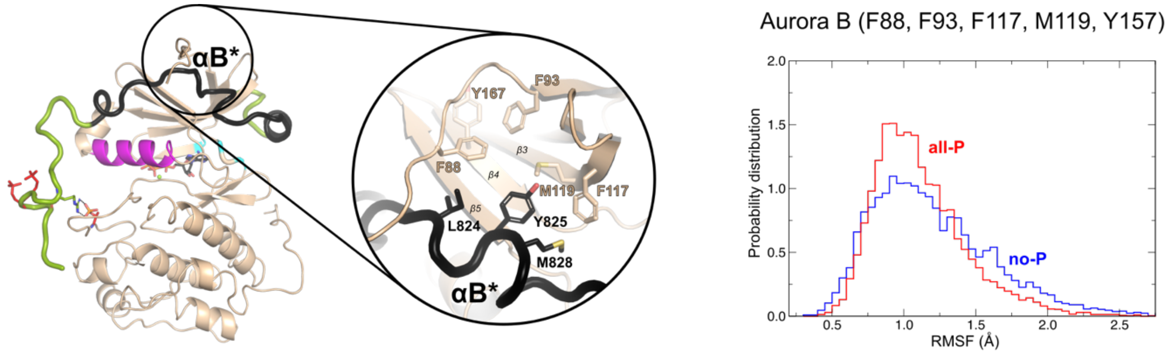


**Fig. S6.**
The ribbon diagram of the [Aurora B/IN-box]$^{all\text{-}P}$ structure (oriented and colored as in Fig. 2). The interface between IN-box$^{\alpha B*}$ and Aurora B is circled and shown in more detail. On the right is the plot of the RMSF for Aurora B (Phe88, Phe93, Phe117, Met119 and Tyr157) with probability distribution. RMSF in [Aurora B/IN-box]$^{all\text{-}P}$ is in red and [Aurora B/IN-box]$^{no\text{-}P}$ is in blue.

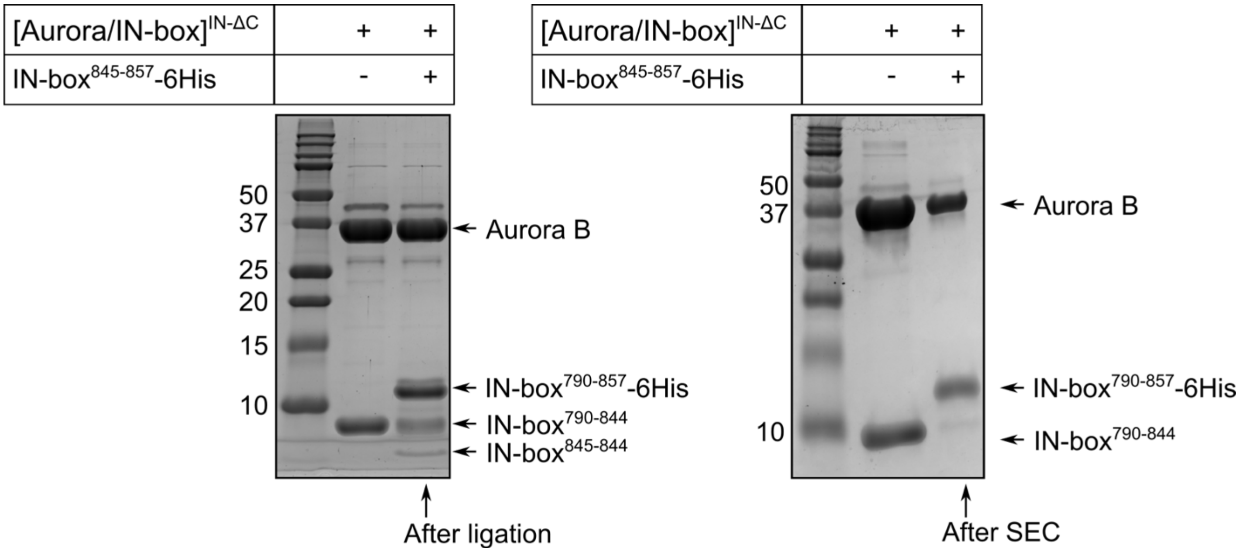


**Fig. S7.**

SDS-PAGE gel showing purification of partially phosphorylated [Aurora B/IN-box] enzymes. The left gel shows the species present in the solution during the ligation process. The right gel shows [Aurora B/IN-box]$^{loop\text{-}P}$ after ligation and size exclusion chromatography. For reference, the first lane on both gels shows the [Aurora B/IN-box]$^{\Delta C}$ complex.

**A.**

Aurora B

$^{248}$pTMCGTLDYLPPEMIEGK$^{264}$

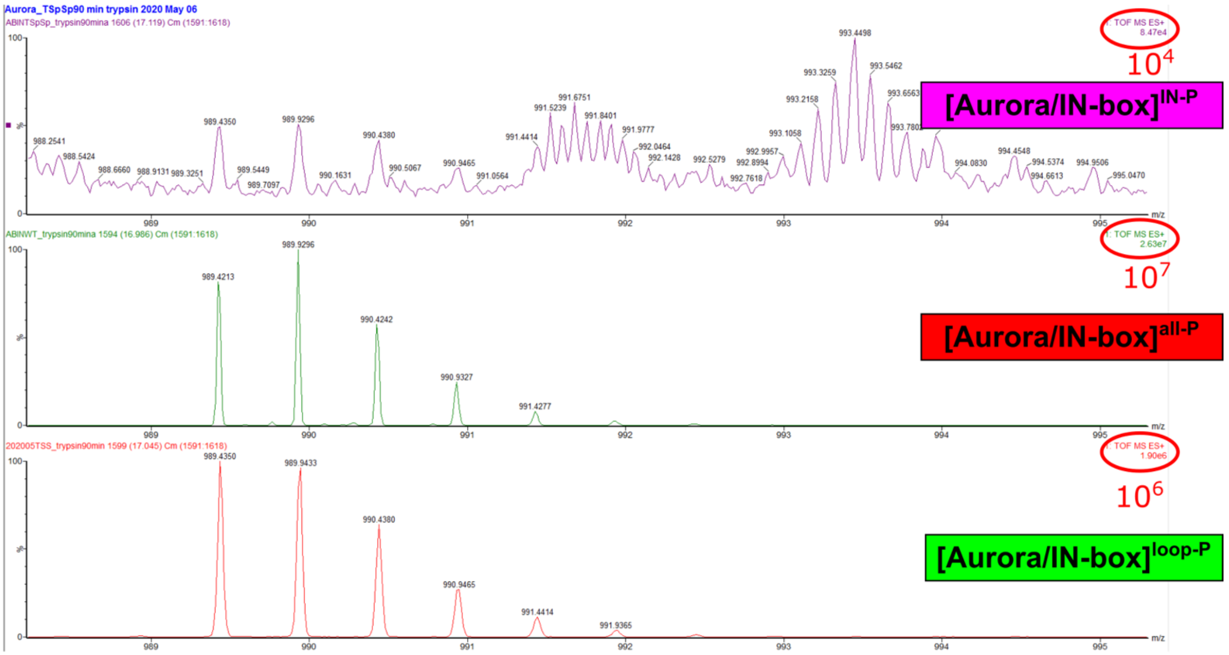


**B.**

IN-box

$^{848}$TpSpSAVWHSPPL$^{858}$-6His

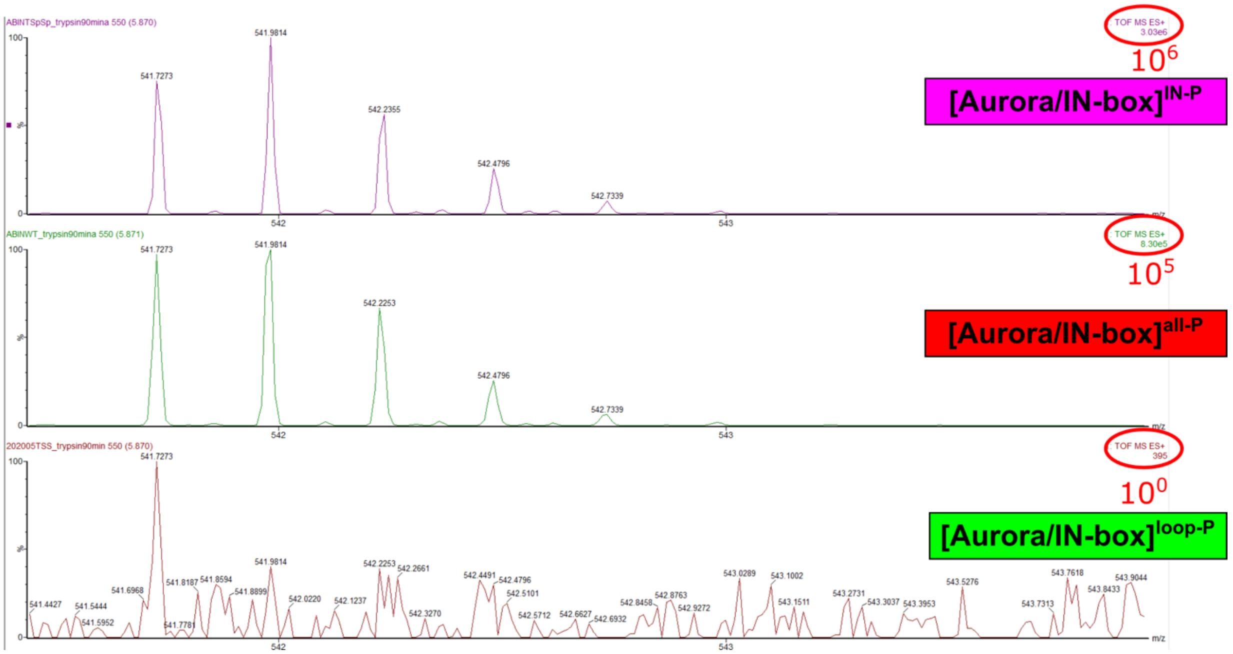

**Fig. S8.**
Mass spectra of peptides containing Aurora $B^{Thr248}$ (A) or IN-$box^{Ser849}$ and IN-$box^{Ser850}$ (B) show the presence or absence of phosphorylation in different [Aurora B/IN-box] complexes.

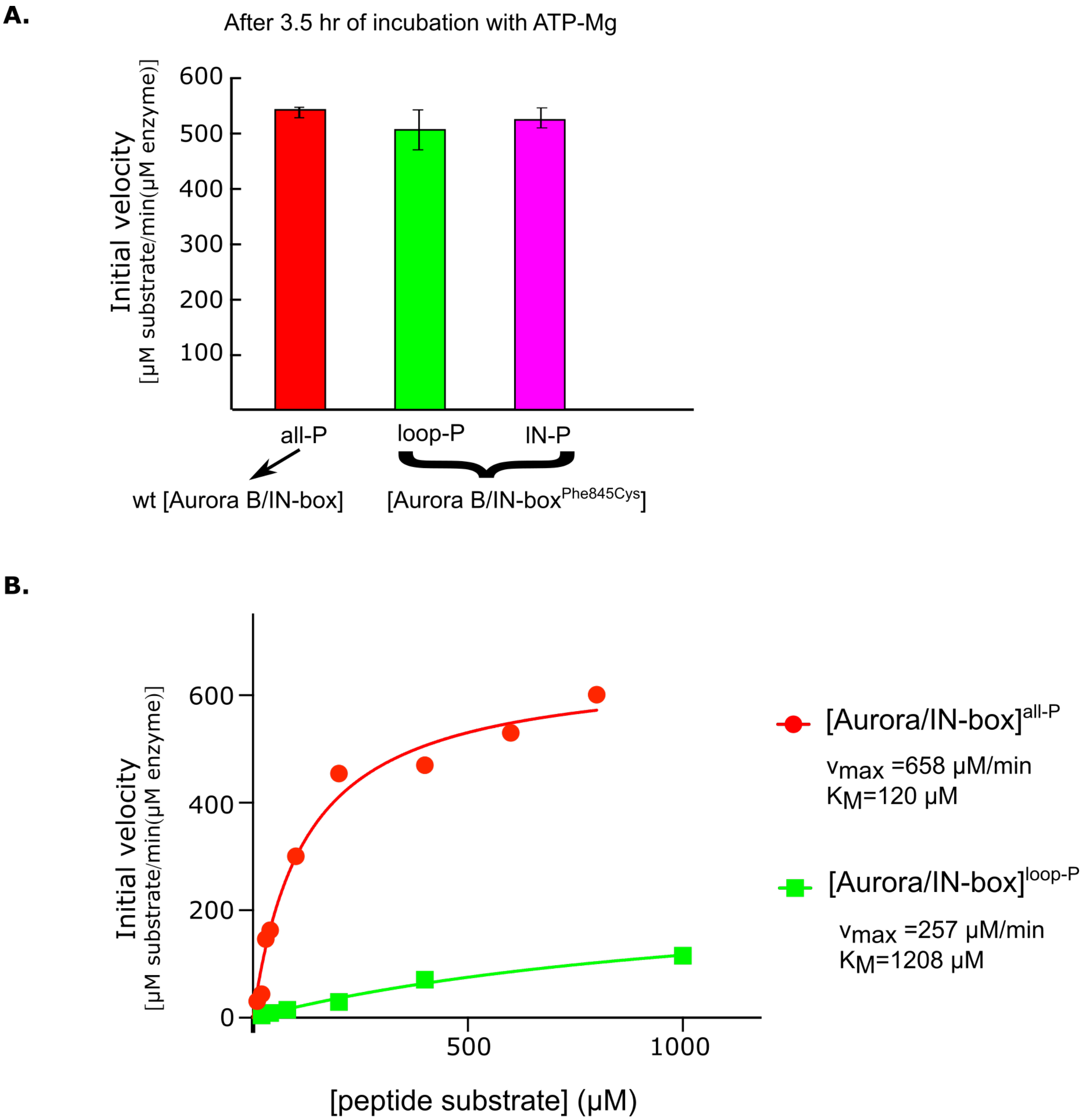


**Fig. S9.**
**A.** After 2.5 h incubation with ATP-Mg, semisynthetic constructs with the IN -box$^{Phe845Cys}$ mutation show the same enzymatic activity as the non-mutated (wt) active [Aurora B/IN-box]. **B.** Dependence of enzyme rate on substrate peptide concentration for [Aurora B/IN-box]$^{all\text{-}P}$ (red) and [Aurora B/ IN -box]$^{loop\text{-}P}$ (green).

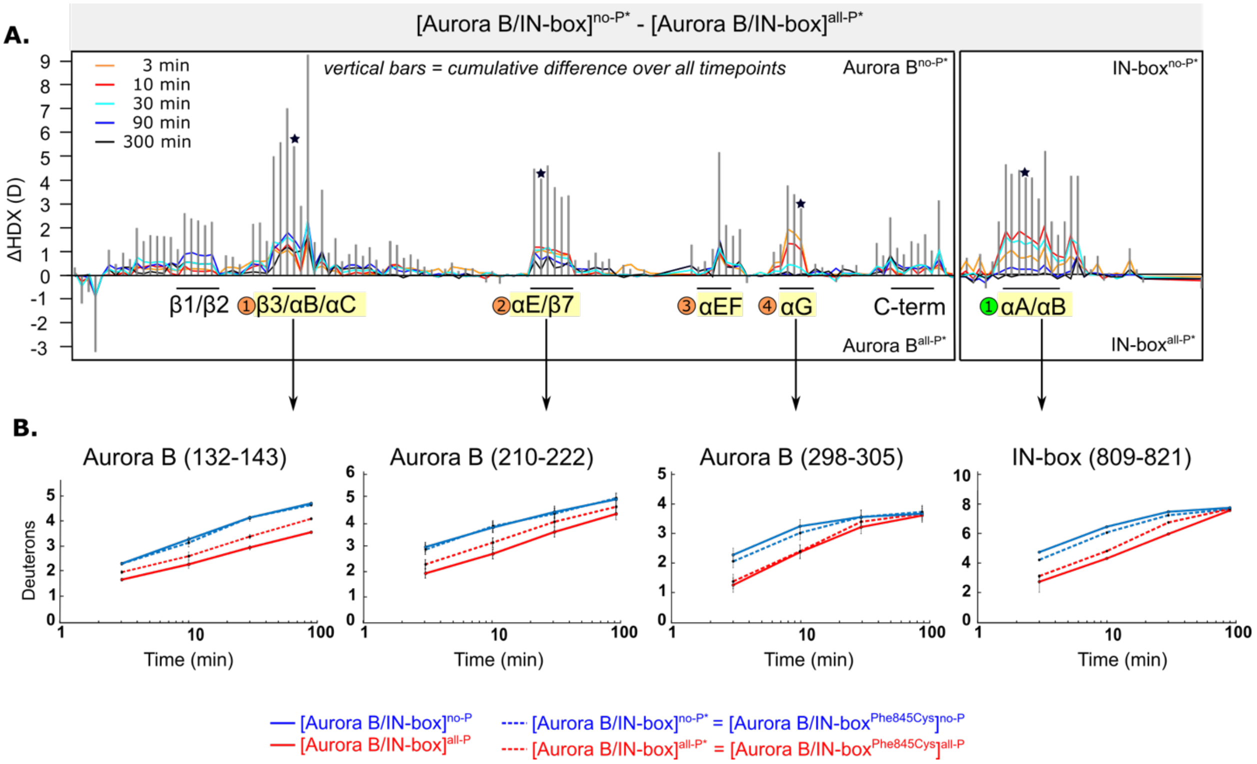


**Fig. S10.**

**A.** Butterfly-difference plot for a comparison of H/D exchange differences between unphosphorylated [Aurora B/IN-box$^{Phe845Cys}$] = [Aurora B/IN-box]$^{no\text{-}P*}$, and fully phosphorylated [Aurora B/IN-box$^{Phe845Cys}$] = [Aurora B/IN-box]$^{all\text{-}P*}$. **B.** Uptake plots for representative peptides (covering the same regions as in Fig. 1 and Fig. 4.) for wt [Aurora B/IN-box] (solid lines) and [Aurora B/IN-box$^{Phe845Cys}$] (dashed lines) in the fully phosphorylated state (red) and unphosphorylated state (blue). Measurements were performed in triplicates, and error bars are shown as black lines for each time point.

#### IN-box
$^{807}$LTQAIRQQYYKPIDV$^{821}$

##### [Aurora/IN-box$^{Phe845Cys}$]$^{no-P}$

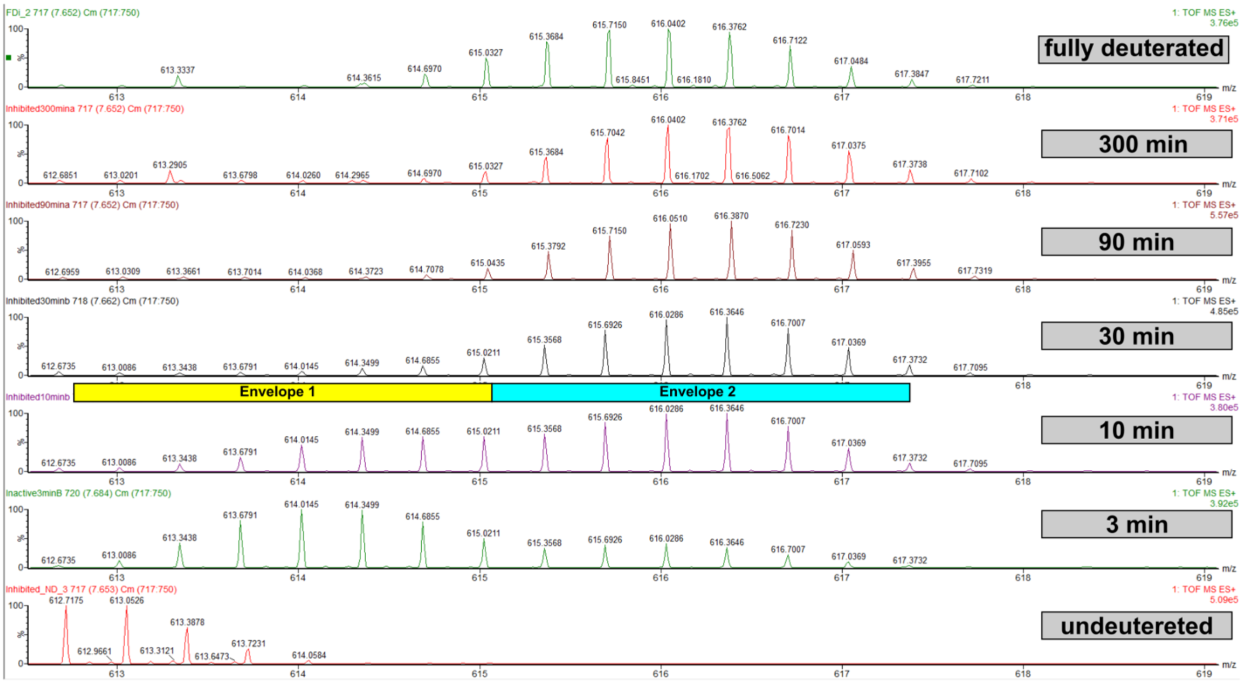


##### [Aurora/IN-box$^{Phe845Cys}$]$^{all-P}$

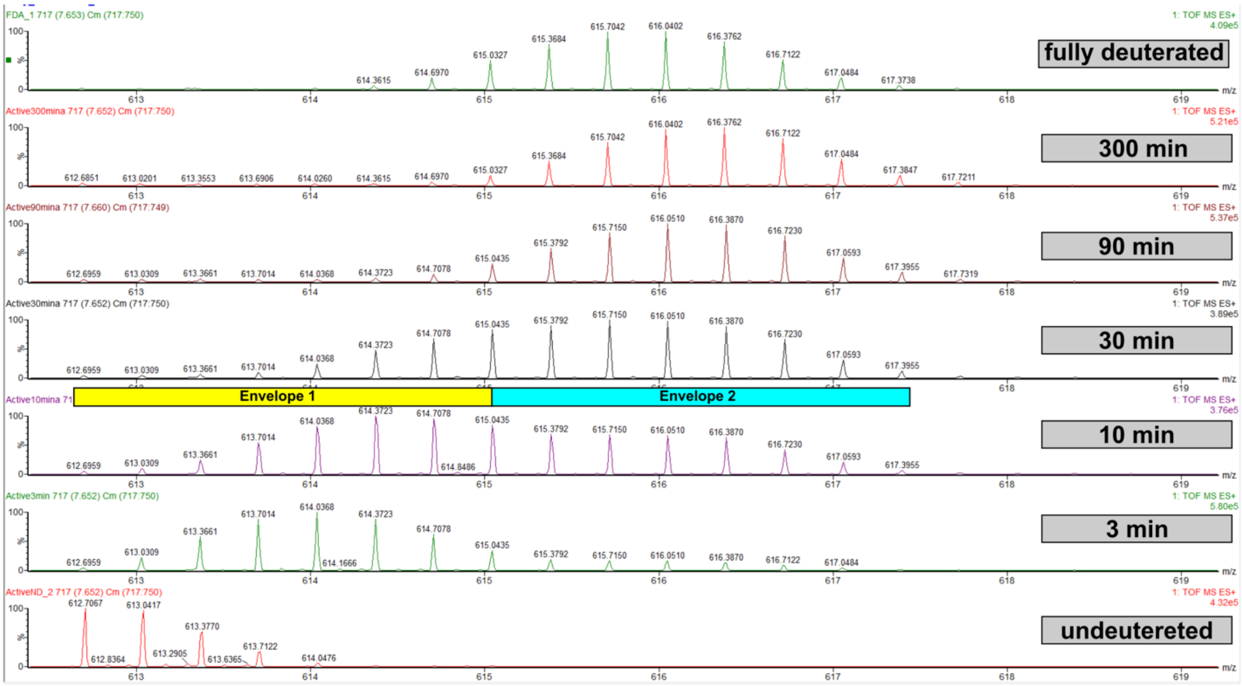

**Fig. S11.**

EX1 kinetics observed in IN-box$^{\alpha A}$, peptide $^{807}$LTQAIRQQYYKPIDV$^{821}$, in [Aurora B/IN-box$^{Phe845Cys}$]$^{no\text{-}P}$ and [Aurora B/IN-box$^{Phe845Cys}$]$^{all\text{-}P}$. Comparable with figure fig. S3.

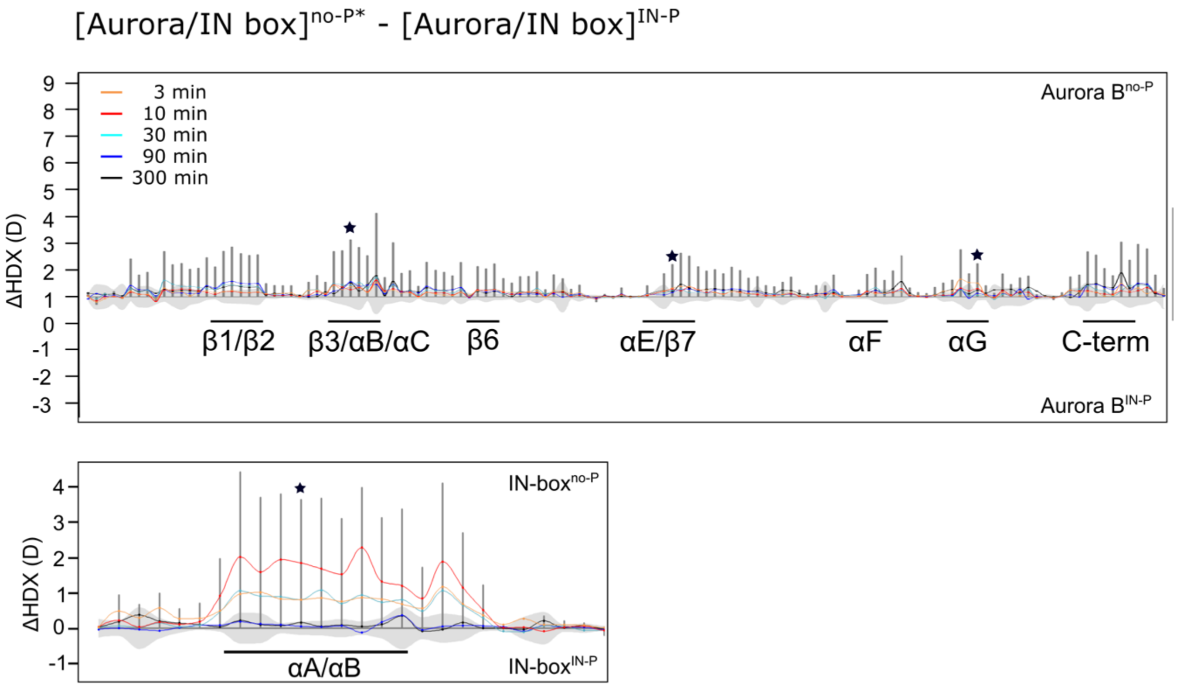


**Fig. S12.**
Difference plot showing the difference in HDX between [Aurora B/IN-box]$^{no-P*}$ and [Aurora B/IN-box]$^{IN-P}$ over 5 time points (orange – 3 min; red - 10 min; cyan - 30 min; blue- 90 min; black – 300 min); as in Figure 1. The asterisk indicates representative peptides for which uptake plots are shown in Figure 4B (purple traces). Note that [Aurora B/IN-box]$^{no-P*}$ carries the same "scar" mutation as [Aurora B/IN-box]$^{IN-P}$ to calibrate for the effects of the mutation on enzyme dynamics.

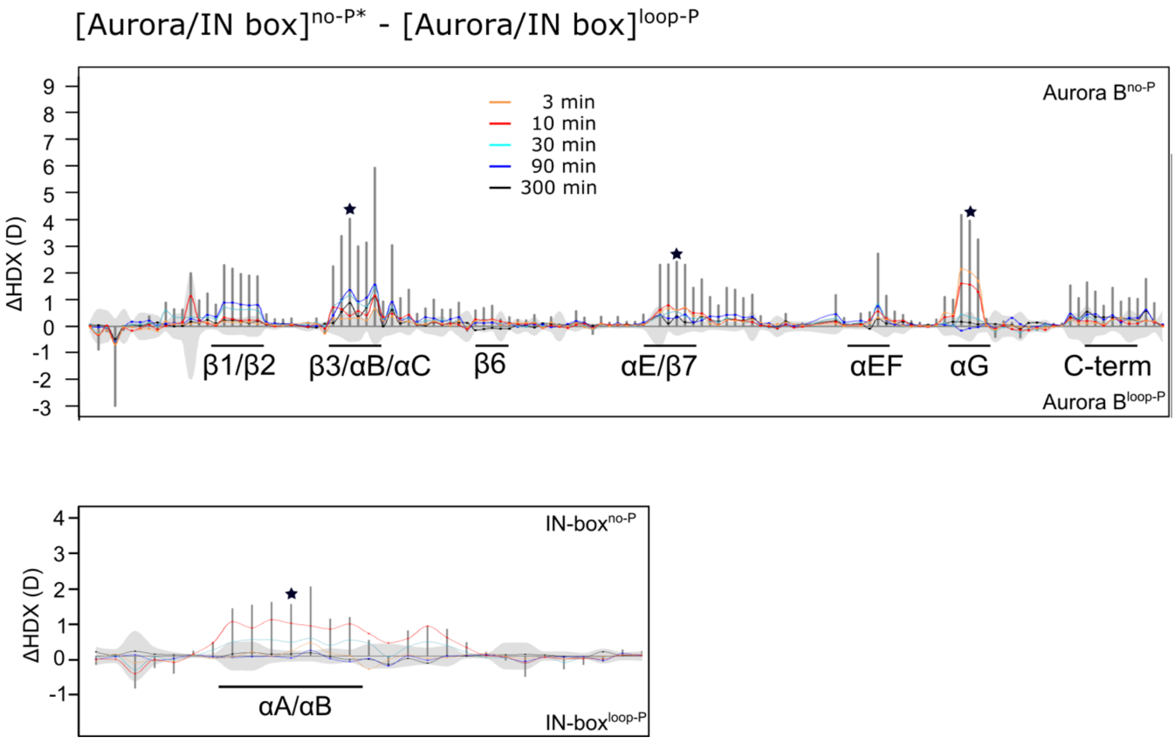


**Fig. S13.**

**A.** Difference plot showing the difference in HDX between [Aurora B/IN-box]$^{no\text{-}P*}$ and [Aurora B/IN-box]$^{loop\text{-}P}$ over 5 time points (orange – 3 min; red - 10 min; cyan - 30 min; blue- 90 min; black – 300 min); as in Figure 1. The asterisk indicates representative peptides for which the uptake plots are shown in Figure 4B (green). Note that [Aurora B/IN-box]$^{no\text{-}P*}$ carries the same "scar" mutation as [Aurora B/IN-box]$^{loop\text{-}P}$ to calibrate for the effects of the mutation on enzyme dynamics.

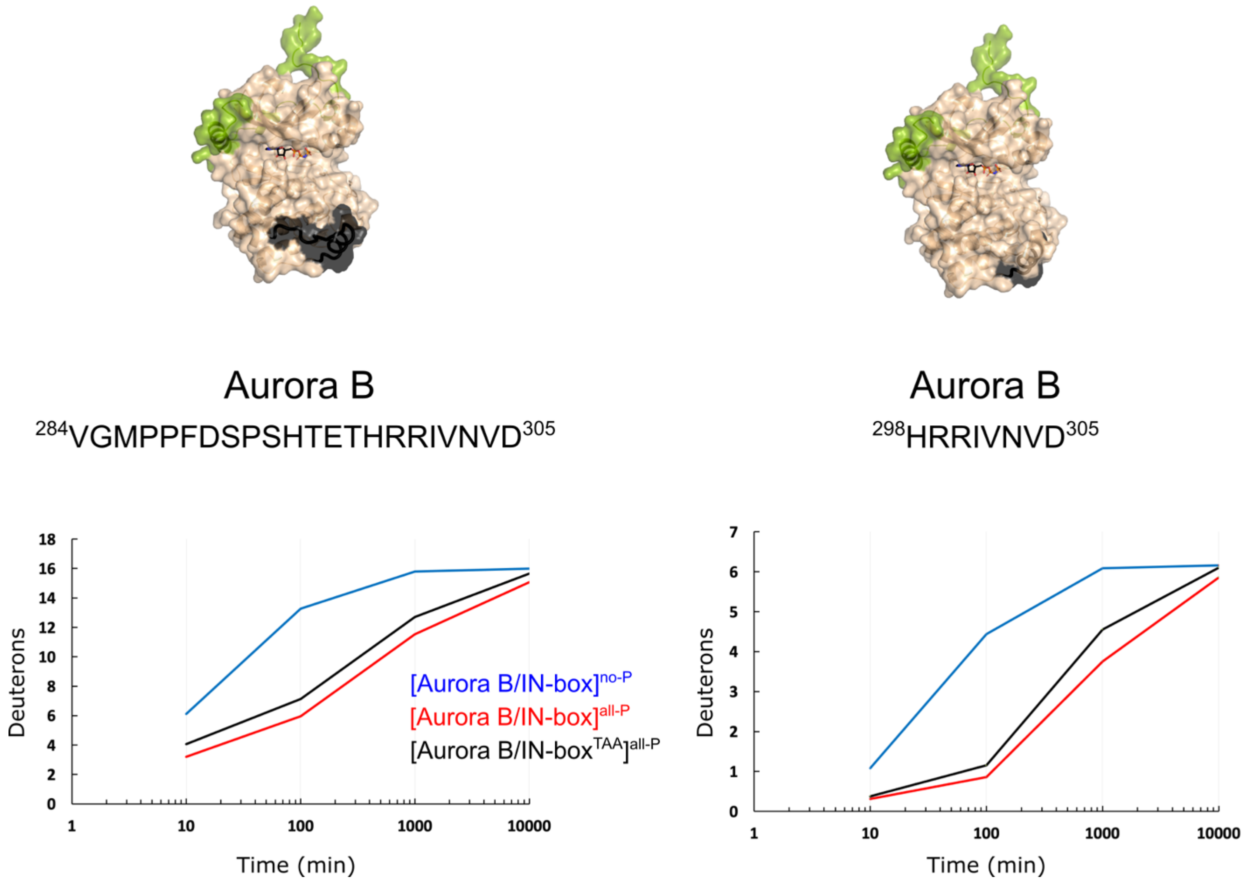


**Fig. S14.**
Structuring of Aurora B$^{\alpha G}$ is independent of phosphorylation in the IN-box TSS motif. The dynamics of the exchange for two peptides covering the Aurora B$^{\alpha G}$ region of the enzyme complex in which the IN-box TSS motif is mutated to TAA so that it cannot be phosphorylated, [Aurora B/IN-box$^{TAA}$], shows a comparable level of protection upon enzyme complex phosphorylation as identical peptides in the phosphorylated wt enzyme complex.

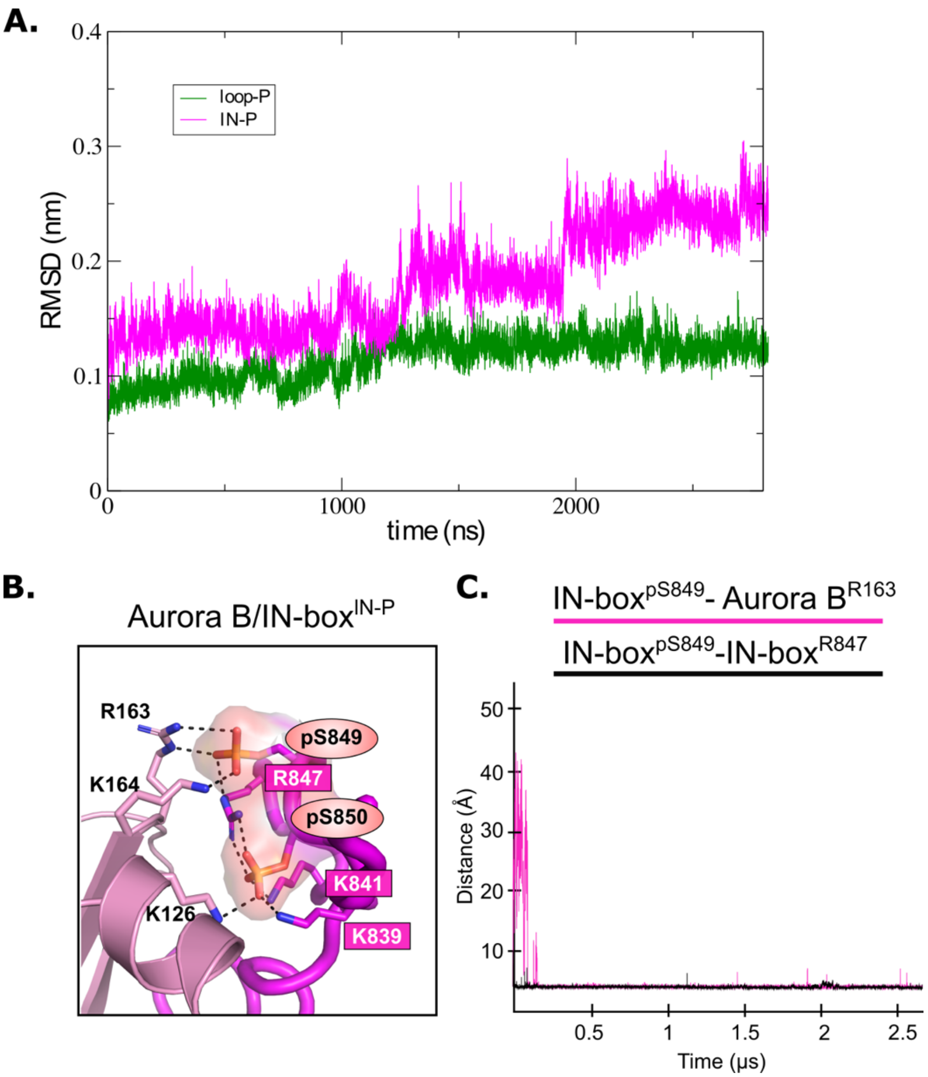


**Fig. S15.**

**A.** Root-mean-square-deviation (RMSD) for [Aurora B/IN-box]$^{\text{IN-P}}$ (magenta) and [Aurora B/IN-box]$^{\text{loop-P}}$ (green) complexes during 2800 ns MD simulations. **B.** Magnified representation of the interactions between the phosphorylated TSS motif (IN-box$^{\text{Ser849-P}}$ and IN-box$^{\text{Ser850-P}}$) and the lysines and arginines of IN-box (dark pink labels) and the N-terminal lobe of Aurora B (left) for [Aurora B/IN-box]$^{\text{IN-P}}$. Right: The change in distance between the phosphorus atom in IN-box$^{\text{pSer249}}$ and the C$\zeta$ atom in Aurora B$^{\text{Arg163}}$ (pink) and the change in distance between the phosphorus atom in IN-box$^{\text{pSer249}}$ and the C$\zeta$ atom in IN-box$^{\text{Arg847}}$ (black) during the MD simulation.

**A.**

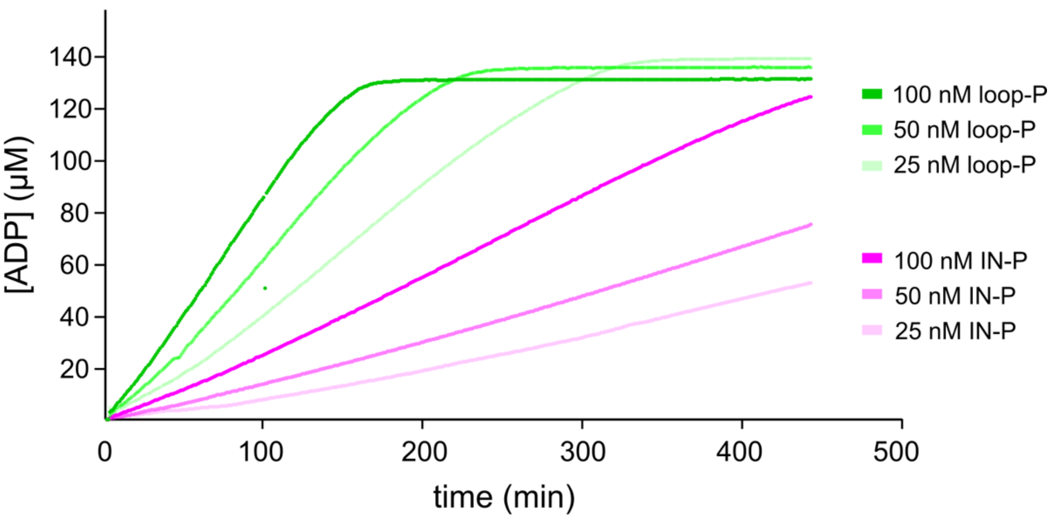


**B.**

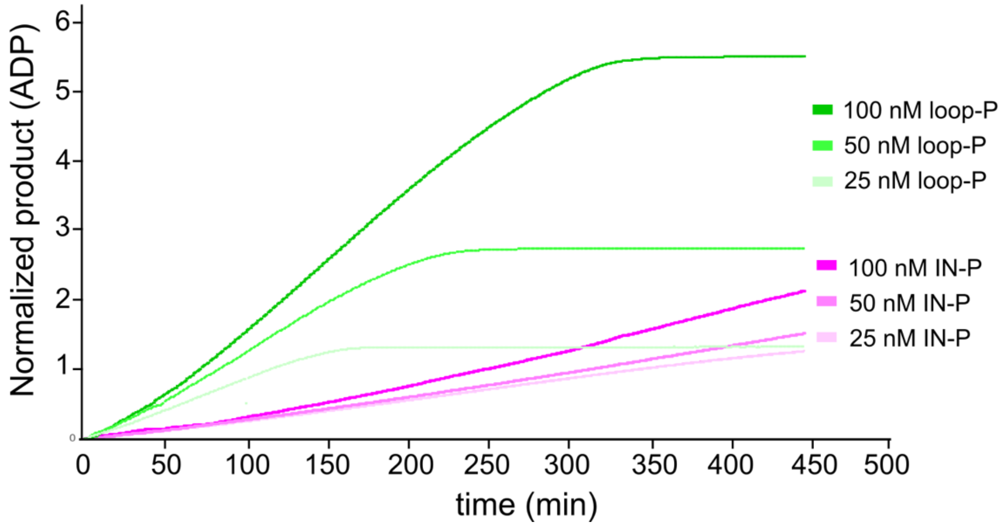


**Fig. S16.**

**A.** Product release curves for [Aurora B/IN-box]$^{loop\text{-}P}$ (green) and [Aurora B/IN-box]$^{IN\text{-}P}$ (magenta) in the presence of the peptide substrate and ATP at different enzyme concentrations (see legend). **B.** The curves from A were normalized to the enzyme concentration. Note that after normalization the pink traces overlap during the first ~50 minutes of incubation, indicating that phosphorylation of the activation loop is an intramolecular, concentration independent, reaction.

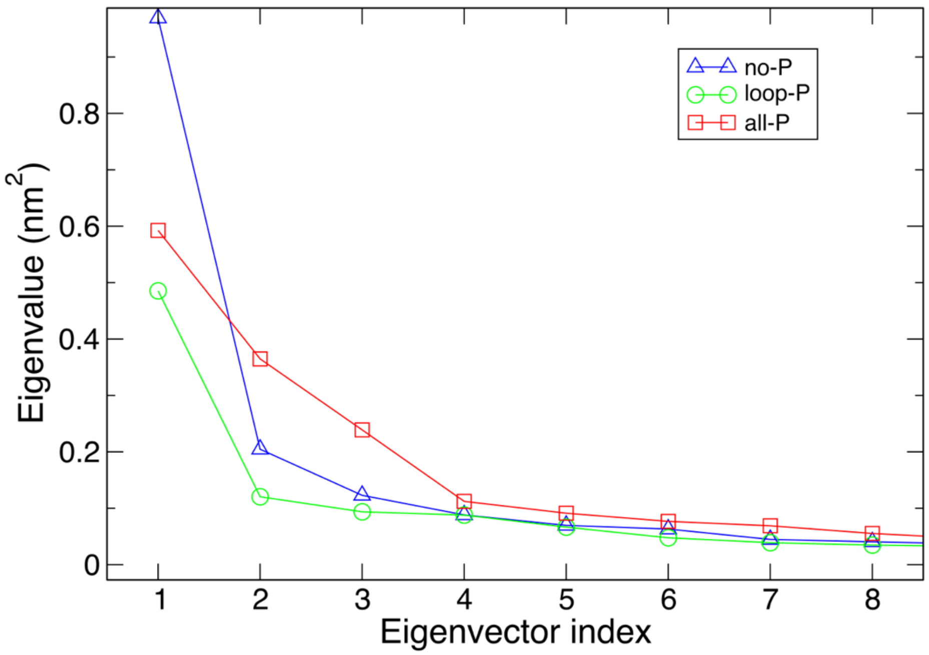


**Fig. S17.**
Eigen values obtained from MD trajectories.

**Data set D1.**

Full HDX data set for [Aurora B/IN-box]$^{all\text{-}P*}$ (red), [Aurora B/IN-box]$^{IN\text{-}P}$ (purple), [Aurora B/IN-box]$^{loop\text{-}P}$ (green) and [Aurora B/IN-box]$^{no\text{-}P*}$ (blue). All collected peptides are shown with the standard error calculated based on two replicates. The black dot in the corner of the graph indicates the exchange of the fully deuterated control for the peptide of interest.

Data set D1 (page 1 of 5)

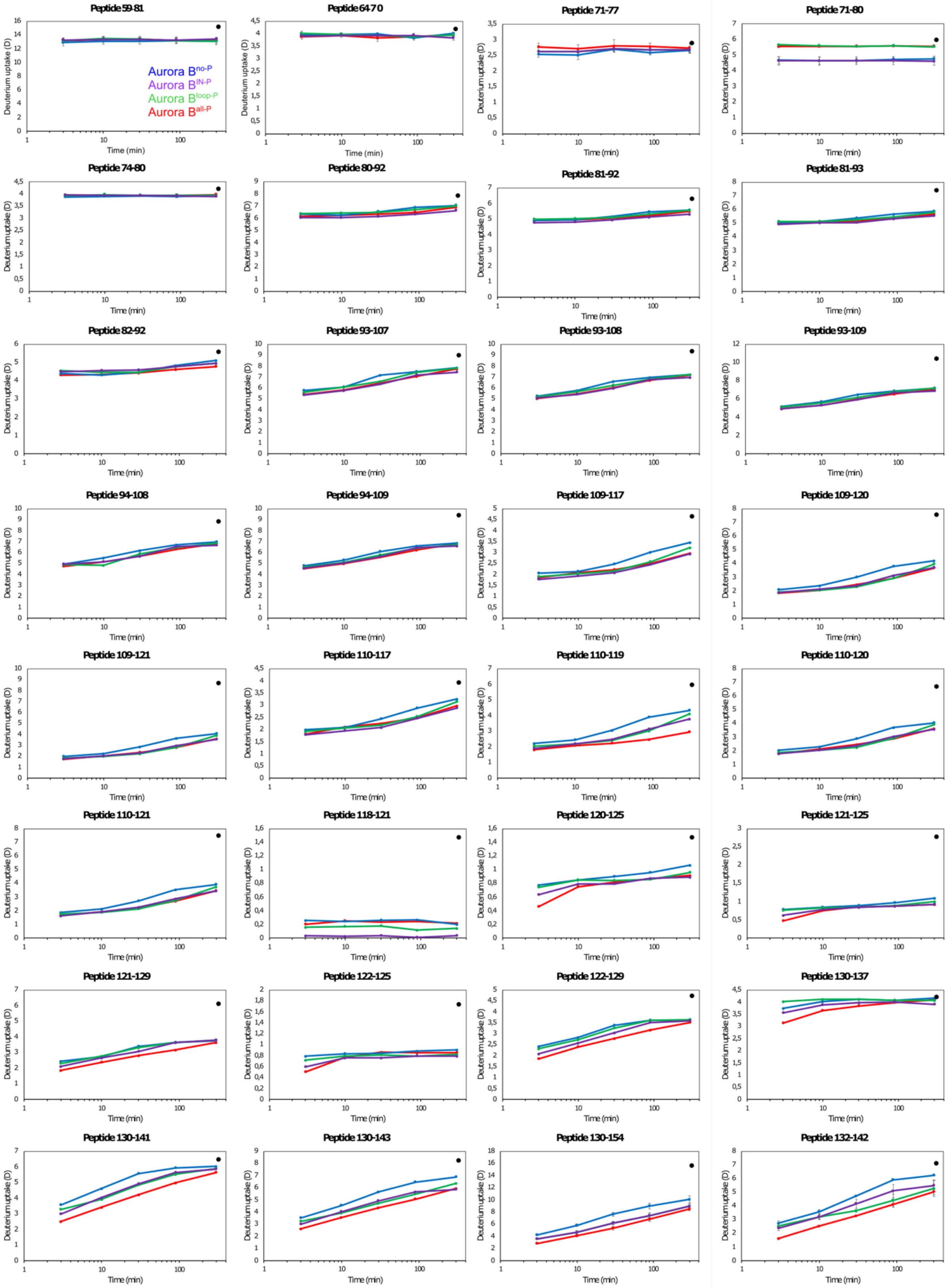

Data set D1 (page 2 of 5)

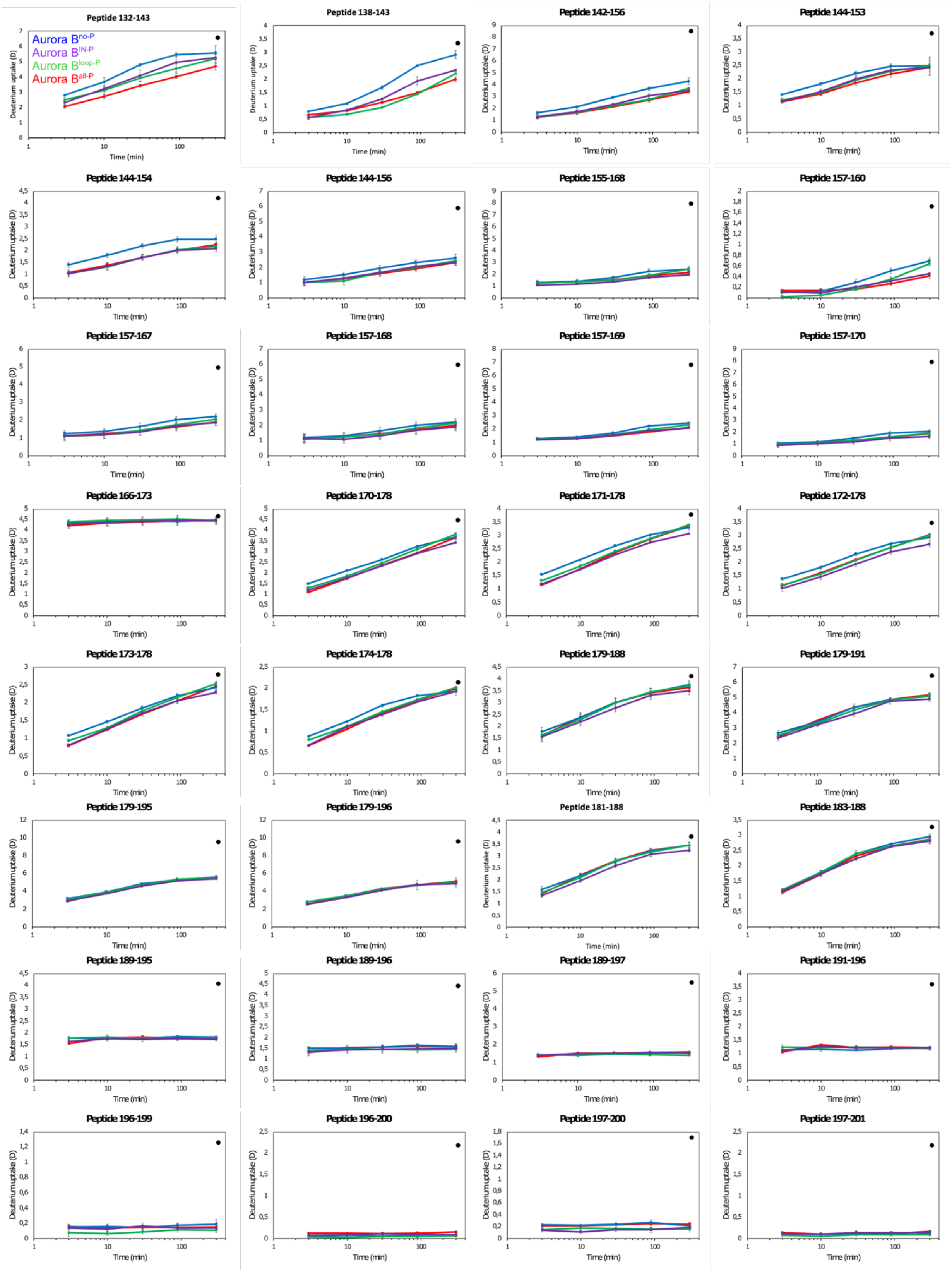

Data set D1 (page 3 of 5)

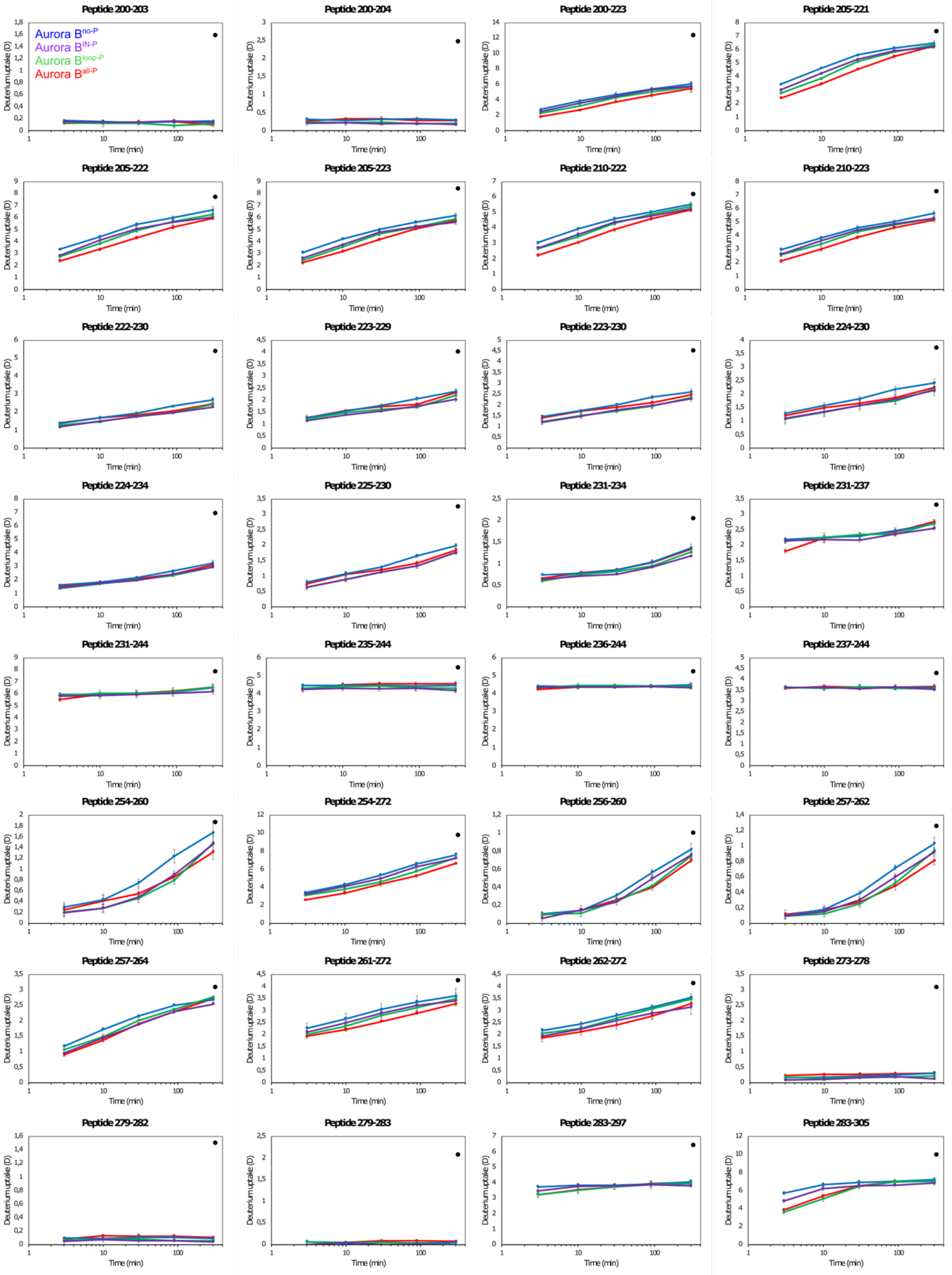

#### Data set D1 (page 5 of 5) **IN-box peptides**

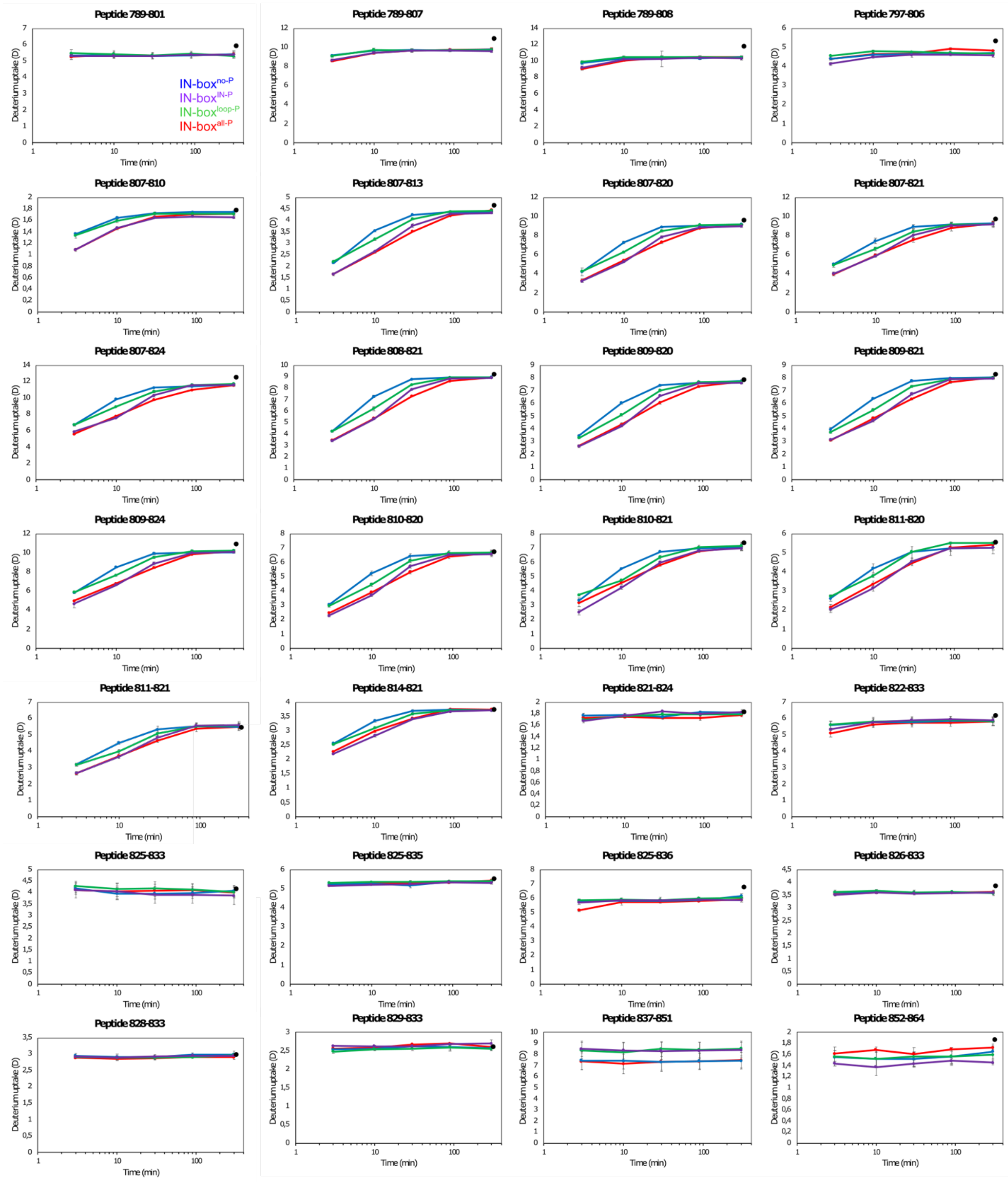

**Movie M1. (Separate file)**
2.8 μs MD simulation trajectory of [Aurora B/IN-box]$^{no\text{-}P}$.

**Movie M2. (Separate file)**
2.8 μs MD simulation trajectory of [Aurora B/IN-box]$^{all\text{-}P}$.

**Movie M3. (Separate file)**
2.8 μs MD simulation trajectory of [Aurora B/IN-box]$^{IN\text{-}P}$.

**Movie M4. (Separate file)**
2.8 μs MD simulation trajectory of [Aurora B/IN-box]$^{loop\text{-}P}$.

**Movie M5. (Separate file)**
PC1 for [Aurora B/IN-box]$^{all\text{-}P}$: Coordinated motions of open-closed/twist and activation loop movements.
